# Principles of cortical interactions in modular recurrent networks

**DOI:** 10.64898/2025.12.19.695176

**Authors:** Deyue Kong, Joe Barreto, Lorenzo Butti, Matthias Kaschube, Benjamin Scholl

## Abstract

Cortical circuits are proposed to amplify weak sensory inputs but transition to feature competition during strong drive. However, the evidence for this is scarce in cortices with a functional, modular organization, such as in primates and carnivores, where neighboring excitatory and inhibitory cells share feature selectivity. Here, we demonstrate that networks in ferret primary visual cortex switch between amplification and competition depending on sensory input. Combining cellular perturbations and statistical modeling, we uncovered broad suppressive influence modulated by visual contrast. At low contrast, functionally-coupled cells exhibited mutual amplification, switching to suppression at high contrast. This reversal emerged in recurrent network models in a Cross-Dominant regime, where inhibitory-excitatory coupling exceeds excitatory-excitatory coupling. These models predicted strong suppression from inhibitory cells, confirmed with cell-type-specific perturbations. Our results provide direct evidence that cortical recurrence with functional, modular organization toggles between amplification and suppression, supporting long-standing predictions from predictions from theory and models of visual cortex.

## Introduction

Cortical circuits must adapt dynamically to the strength of sensory inputs, integrating information when signals are weak and refining representations when they are strong. Experimental and theoretical studies suggest this adaptation requires recurrent cortical circuits that engage in cooperative amplification and competitive interactions within functional networks^1–5^. Recent studies in mouse primary visual cortex (V1) have begun to support this idea^6–10^, and the observations from this experimental work have been explained by the emergence of novel circuit motifs whereby excitatory and inhibitory cells are strongly connected^11–13^. Despite these advancements, neither models nor data provide a clear picture of the behavior of cortical columns within circuits with a functional modular organization, such as the visual cortex of carnivores and primates. Within these circuits, neighboring cells (excitatory and inhibitory) are selective for similar features in visual space^14^, raising the question as to whether cortical columns act as amplifiers or engage in feature competition^15^. Moreover, columnar network models make a strong prediction: cortical circuits should engage in functional amplification during weak sensory drive, transitioning to suppressive interactions with increasing sensory drive^2,16,17^-- a prediction which has yet to be directly tested experimentally.

Here we used single-cell perturbations in ferret V1 to show that cortical networks switch between amplification and suppression, depending on stimulus strength. We combined two-photon optogenetic stimulation with a generalized linear model (GLM) to quantify perturbations, revealing spatially broad suppressive influence modulated by stimulus contrast. At high contrast, we observed suppression, and at low contrast these networks switched to amplification, but only between functionally-coupled cells. This reversal emerged in a recurrent network model with strong inhibitory-to-excitatory connectivity and weaker excitatory-to-excitatory connections. In addition, we were able to capture most features observed experimentally with a reduced and analytically tractable recurrent model. Our mathematical analyses predict markedly stronger suppressive influences from inhibitory cells onto excitatory populations, which we confirmed with cell-type-specific perturbations. Overall, our results show that cortical recurrence with modular networks toggles between amplification and suppression, providing direct evidence long-standing predictions from theories and circuit models of visual cortex.

## Results

To measure the functional influence of single excitatory cells in cortical circuits with a modular organization, we performed two-photon photostimulation of single cells during two-photon population calcium imaging in layer 2/3 ferret V1 *in vivo* (**Fig. 1**). We co-expressed the calcium sensor jGCaMP8s^18^ and a red-shifted soma-targeted channelrhodopsin optimized for two-photon excitation (ST-ChroME)^19^. Both constructs were driven under a hSyn promoter for selective targeting of excitatory cells (see Methods)^14^. We imaged the calcium activity of all labeled cells in a given field of view (FOV, ∼750 x 750 µm) while presenting drifting gratings varying in direction (0 - 315 deg) and contrast (8 - 100%) (see Methods), targeting single cells and sham locations (e.g. blood vessels, neuropil) for photostimulation (**Fig. 1a-c**). Within single FOVs, visually-driven activity during photostimulation exhibited a classic columnar organization of direction/orientation preference^14,20^ (**Fig. 1d**).

**Figure 1.**
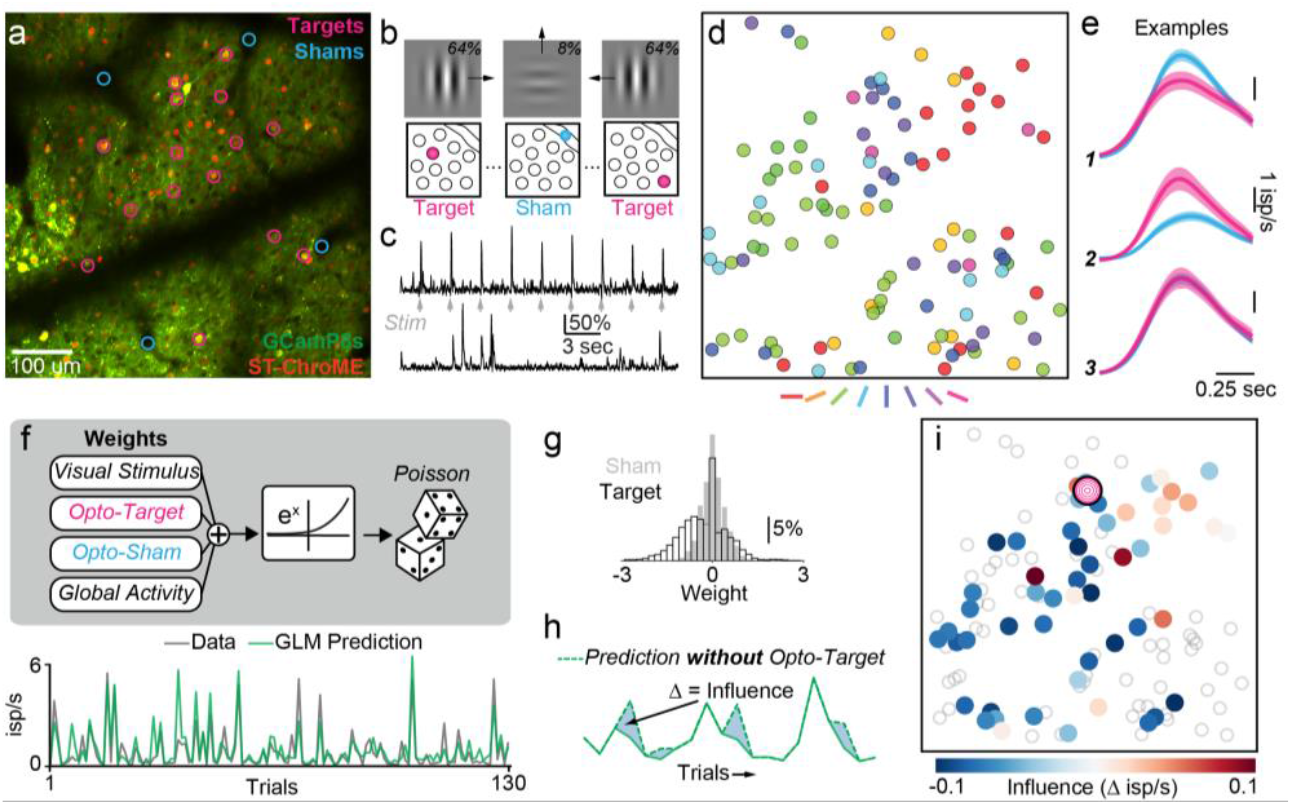
Inferring single-cell perturbation influence between excitatory cells in ferret visual cortex. (a) Example field-of-view (FOV) with target (pink) and sham (blue) locations marked. Image is a merge of the average projection (1000 frames) of green and red PMT channels. All cells expressing GCaMP8s and ST-ChroME are excitatory. (b) Schematic of experimental paradigm. Drifting grating stimuli varying in direction and contrast were presented randomly and on each trial a target or sham location was photostimulated. (c) Calcium activity from a target cell when photostimulated during spontaneous activity and neighboring non-targeted cell. (d) Columnar organization of orientation preferences for the cells in the FOV in **a.** (e) Average responses for three example nontarget cells for sham trials (blue) and trials when a single target cell was photostimulated (pink). Responses averaged over all trials and stimulus conditions. (f) Generalized linear model (GLM) used to predict trial-by-trial activity (inferred spikes/sec) of nontarget cells. Example data (gray) and model prediction (green) for a subset of trials is shown below. (g) Sham GLM weights are used to both constrain target GLM weights and provide a significance test for each target-nontarget pair (see Methods). (h) Influence is calculated by taking the difference between GLM predictions with and without that target’s regressor weight. (i) Example influence map for a single photostimulated target cell (pink spiral), for the FOV in **a**. Gray circles indicate nontarget cells not significantly influenced by the target.

Single-cell perturbations were conducted with galvo-galvo spiral scanning of a high-repetition rate infrared laser (1070 nm, 80 Mhz) driven by a Bruker 2P Plus system with built-in spatial calibration routines. Photostimulation protocol followed Chettih and Harvey (2018), with a total duration of 250-500 ms (see Methods). In each FOV, we first targeted several cells to measure laser power response curves during spontaneous activity (**Supplementary Figure 1**). Average power for photostimulation during visual stimulation was chosen to be the smallest value to drive calcium responses before saturation, reliably activating cells in a spatially-precise manner (**Supplementary Figure 1**). Given cortical dynamics following the onset of visual stimuli, we aimed to perturb cells during the steady-state response period by triggering photostimulation ∼120 ms after stimulus presentation (and subsequent feedforward input to layer 2/3 V1^21^).

We expected weak influences from perturbing single excitatory cells^6,11,13,19,22–24^ (examples shown in **Fig. 1e**), so to quantitatively estimate the potential influence of a target onto neighboring nontarget cells we used a generalized linear model (GLM)^25^ (see Methods) to predict trial-to-trial activity, measured as the peak change in spike rate inferred from calcium ΔF/ F transients^26^ (**Fig. 1f**). This allowed us to isolate the effects of single-cell perturbation from sensory-driven activity and global correlations, which generally account for a majority of variance in trial-to-trial activity^25^. Target photostimulation and sham weights were fit for each visual stimulus condition. Significant weights were identified by comparing influence weights for individual target cells with the distribution of sham weights for a given FOV (**Fig. 1g**). Finally, we defined the (stimulus-dependent) ‘influence’ of a target cell onto a nontarget as the trial-averaged difference in predictions from the full GLM and the same model with the target photostimulation regressor nullified (**Fig. 1h**). Using this metric, we estimated the influence of a target cell onto surrounding neighbors for significant interactions and individual stimulus conditions, generating influence maps across cortical space (**Fig. 1i**). With this approach, we measured influences between 183 target and 2,312 nontarget cells within excitatory networks of ferret V1 across 23 FOVs (n = 8 animals). On average, GLMs explained about half of response variance on withheld trials (train = 56% ± 7%, test = 49% ± 9%, mean ± s.d.).

Theoretical studies suggest that strong local excitation and lateral inhibition (LELI), sometimes referred to as a ‘Mexican-Hat’ connectivity profile, underlies functional interactions in a columnar cortex such as ferret V1^27–30^. Neighboring cells with similar preferred orientation exhibit facilitatory interactions, becoming suppressive as the cortical distance and tuning difference between target and nontarget cells increase (**Fig. 2a**). Such a profile, albeit on a more local scale, has been observed in mouse V1^6,7,24^, raising the possibility that excitatory influences in ferret V1 exhibit a more pronounced and spatially widespread LELI profile. To test this prediction, we examined influence as a function of cortical distance (**Fig. 2b**), excluding comparisons within 20 µm of photostimulation targets. Under both low and high stimulus contrast, net influence is primarily suppressive. Although suppression is weakest for nearby target cells (< ∼100 - 200 µm), this profile is inconsistent with a columnar-scale LELI profile, where we would expect a clear minimum around 400 µm or about half the typical distance between neighboring iso-orientation column centers (**Supplementary Fig. 2**). Moreover, under low contrast stimulation, excitatory influences are significantly less suppressive both locally and at long-range. Contrast-dependent modulation of suppression is specifically driven by target-nontarget excitatory pairs with similar orientation tuning preference (**Fig. 2c**) and abolished for pairs with orthogonal tuning (**Fig. 2d**). These findings hold when only considering nontarget cells with GLM’s capturing trial-to-trial responses (see Methods; **Supplementary Fig. 2**).

**Figure 2.**
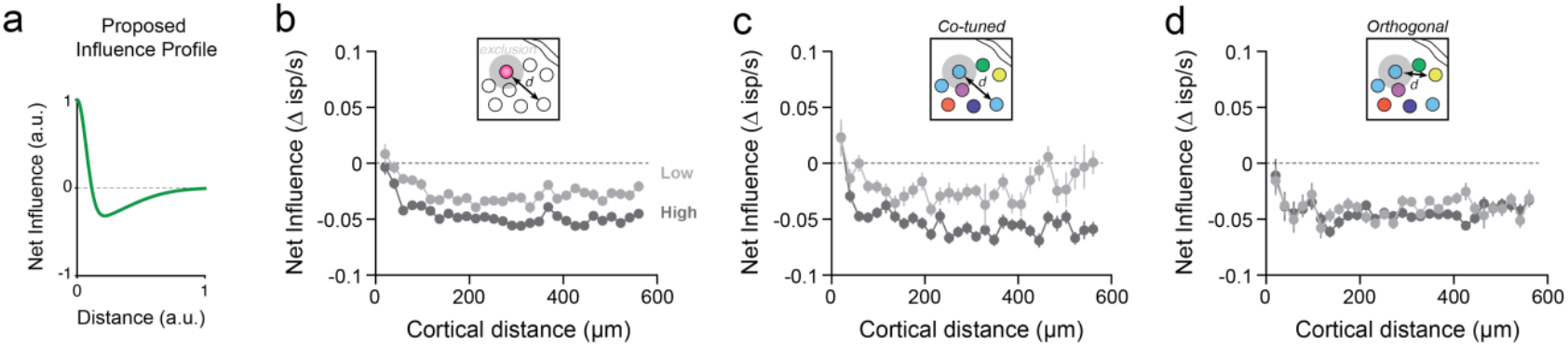
Single excitatory cells exhibit broad suppressive influence, modulated by visual stimulus contrast. (a) Hypothesized LELI (local excitation and lateral inhibition) influence profile: local amplification and longer-range competition. In mouse V1, influences extend a short distance (< 200 µm) ^6,7,24^. (b) Cortical distance-dependent net influences for high (dark gray) and low (light gray) contrast stimuli. Influence is measured in units of change in inferred spikes per second. Distances less than 20 µm were excluded for potential photostimulation artifacts. Each data point represents mean and standard error for that distance bin. Local (20-200 µm) and long-distance influences (>200 µm) were more suppressive at high contrast (p < 0.001 for both, two-sample t-test with unequal variance). In addition, across all distances, low contrast influences were larger than high contrast (low mean = = -0.028 ±0.16 s.d.; high mean = -0.048 ±0.12 s.d.; p < 0.001, two-sample t-test with unequal variance). Data shown are 48,649 and 44,093 comparisons for high and low contrast stimuli, respectively. (c) Same as in **b** for co-tuned excitatory cells (orientation tuning preference difference < 30 deg). Local (20-200 µm) and long-distance influences (>200 µm) were more suppressive at high contrast (p < 0.001 for both, two-sample t-test with unequal variance). Data shown are 11,758 and 10,755 comparisons for high and low contrast stimuli, respectively. (d) Same as in **b** for orthogonally-tuned excitatory cells (orientation tuning preference difference > 30 deg). Local influences were not significantly different (p = 0.051, two-sample t-test with unequal variance). At long distances, low contrast influences were slightly larger than high contrast (low mean = = -0.040 ±0.14 s.d.; high mean = -0.044 ±0.12 s.d.; p = 0.006, two-sample t-test with unequal variance). Data shown are 14,773 and 13,854 comparisons for high and low contrast stimuli, respectively.

As low contrast stimuli typically drive smaller responses in cortical cells, it is possible that contrast-dependent influence onto a cell depends on its response amplitude, rather than network interactions during sensory stimulation. We examined this possibility by comparing distributions of low and high contrast influence conditioned on a nontarget cell’s visual response amplitude on sham trials. Comparing influences measured for the weakest responses during high contrast stimuli (bottom 25%) with low contrast influences, we still observe that low contrast stimuli drive less suppression (**Supplementary Figure 3**). The same is found when comparing high contrast responses with the largest amplitude (top 25%) low contrast responses (**Supplementary Figure 3**). Under both stimulus conditions, we observe that visual response amplitude of nontarget cells is weakly correlated with influence during target stimulation.

Cortical connectivity and perturbation-driven influences are proposed to depend on the functional similarity or functional coupling between cells^6–8,23,31^. In ferret V1, cells are organized in columns representing a common stimulus feature in specific locations of visual space^14,32^. Thus, we next examined how influence between photostimulation target and nontarget excitatory cell depends on functional coupling. Noise correlations are a common measure of functional coupling between cell pairs, introducing less bias than tuning similarity or signal correlations (see Methods; **Fig. 3a**). Noise correlations depend on cortical distance and functional similarity between cells in primate V1^33^ and in mouse V1. Correlated fluctuations are also predictors of monosynaptic connections^34^. As expected, noise correlations in our data decrease with cortical distance (reaching baseline at ∼300 µm), similar to orientation preference difference measured during sham trials and GLM tuning weight correlation (**Supplementary Fig. 4**). Noise correlations co-vary with GLM tuning similarity and orientation preference difference (**Supplementary Fig. 4, Fig. 3a**).

**Figure 3.**
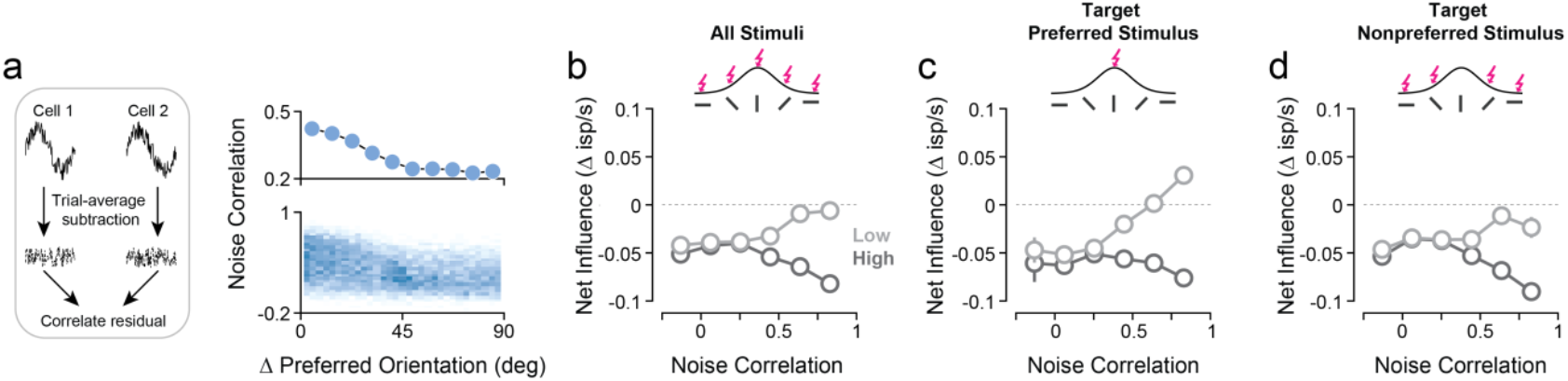
Excitatory influence exhibits amplification and suppression between functionally-coupled cells. (a) Relationship between noise correlation and orientation tuning preference. Noise correlations calculated as correlation of mean-subtracted responses for each stimulus condition (residuals) between cell pairs (*left*). Correlated pairs share similar orientation preference (*right*; circular-linear correlation = 0.39, p < 0.001), but high correlations are not exclusively found between co-tuned pairs. (b) Relationship between local net influence and trial-by-trial correlations for low (light gray) and high (dark gray) contrast. Data points are mean and standard error across all photostimulation trials. Spearman’s *r*_*high contrast*_ = -0.090, n = 45,816 comparisons, p < 0.001. Spearman’s *r*_*low contrast*_ = +0.14, n = 47,845 comparisons, p < 0.001. Target-nontarget pairs with high noise correlation (r > 0.75) exhibited less suppression for low contrast stimuli (low: mean = -0.006 ±0.13 s.d., high: mean = -0.080 ±0.10 s.d., p < 0.001). (c) Same as in **b** for photostimulation trials aligned with the target cell’s preferred orientation (< 30 deg). Spearman’s *r*_*high contrast*_ = -0.005, n = 7,135 comparisons, p = 0.79. Spearman’s *r*_*low contrast*_ = +0.24, n = 7,402 comparisons, p < 0.001. Target-nontarget pairs with high noise correlation exhibited facilitatory influence at low contrast stimuli (low: mean = +0.036 ±0.06 s.d., high: mean = -0.070 ±0.10 s.d., p < 0.001). (d) Same as in **b** for photostimulation trials *not* aligned with the target cell’s preferred orientation (> 30 deg). Spearman’s *r*_*high contrast*_ = -0.11, n = 31,422 comparisons, p = < 0.001. Spearman’s *r*_*low contrast*_ = +0.11, n = 33,012 comparisons, p < 0.001. Target-nontarget pairs with high noise correlation exhibited less suppression for low contrast stimuli (low: mean = -0.026 ±0.15 s.d., high: mean = -0.091 ±0.11 s.d., p < 0.001).

Influence between excitatory cells strongly depends on functional coupling and stimulus contrast (**Fig. 3b**). At low contrast, we observe a trend towards facilitatory interactions between highly correlated cells, but these interactions invert at high contrast, showing stronger suppression. This contrast-dependent switch is also apparent when examining trial-by-trial signal correlations, and to a much lesser degree for similarity in orientation tuning preference (**Supplementary Fig. 5**). We reasoned that for enhancing the detection of weak stimuli, such a switch should make influences more positive primarily by target cells actually driven by the visual stimulus, and so we examined interactions when perturbation trials matched a given target’s preference. Indeed, at low contrast, we observe an even stronger positive relationship between net influence and functional coupling, with suppression during high contrast stimuli (**Fig. 3c**), indicating that the switching effect is highly stimulus-specific. For mismatched perturbation trials we still observe a contrast-dependent switch, but with more suppressive influences (**Fig. 3d**), suggesting competitive interactions are engaged when perturbing a target cell during visual stimuli which would not normally drive it.

How might ferret V1 circuits give rise to pervasive suppressive interactions during excitatory perturbation? To address this question, we explored a rate-based recurrent network consisting of excitatory and inhibitory nonlinear units. Connections between units are defined by a spatial Gaussian profile, emulating a columnar architecture. Similar to experiments and to theoretical work in mouse V1^12,13^, we perturbed the activity of a single excitatory unit around the steady-state response (i.e. fixed point) to a constant stimulus and calculated a targeted unit’s influence (see Methods; **Fig 4a**). Given the small influence of single-cell perturbations, we linearized the network dynamics around this fixed point to measure influence as a change in firing rate. We first considered a classic model of ferret V1: LELI connectivity under conditions which also generate columnar/modular activity patterns (see Methods)^27,30^. For simplicity, we fixed the spatial extent of connectivity (*σ*_*I*_ /*σ*_*E*_ = 1.5). In contrast to our experimental data, single-cell influences within this network showed strong, local amplification (**Fig. 4b**).

**Figure 4.**
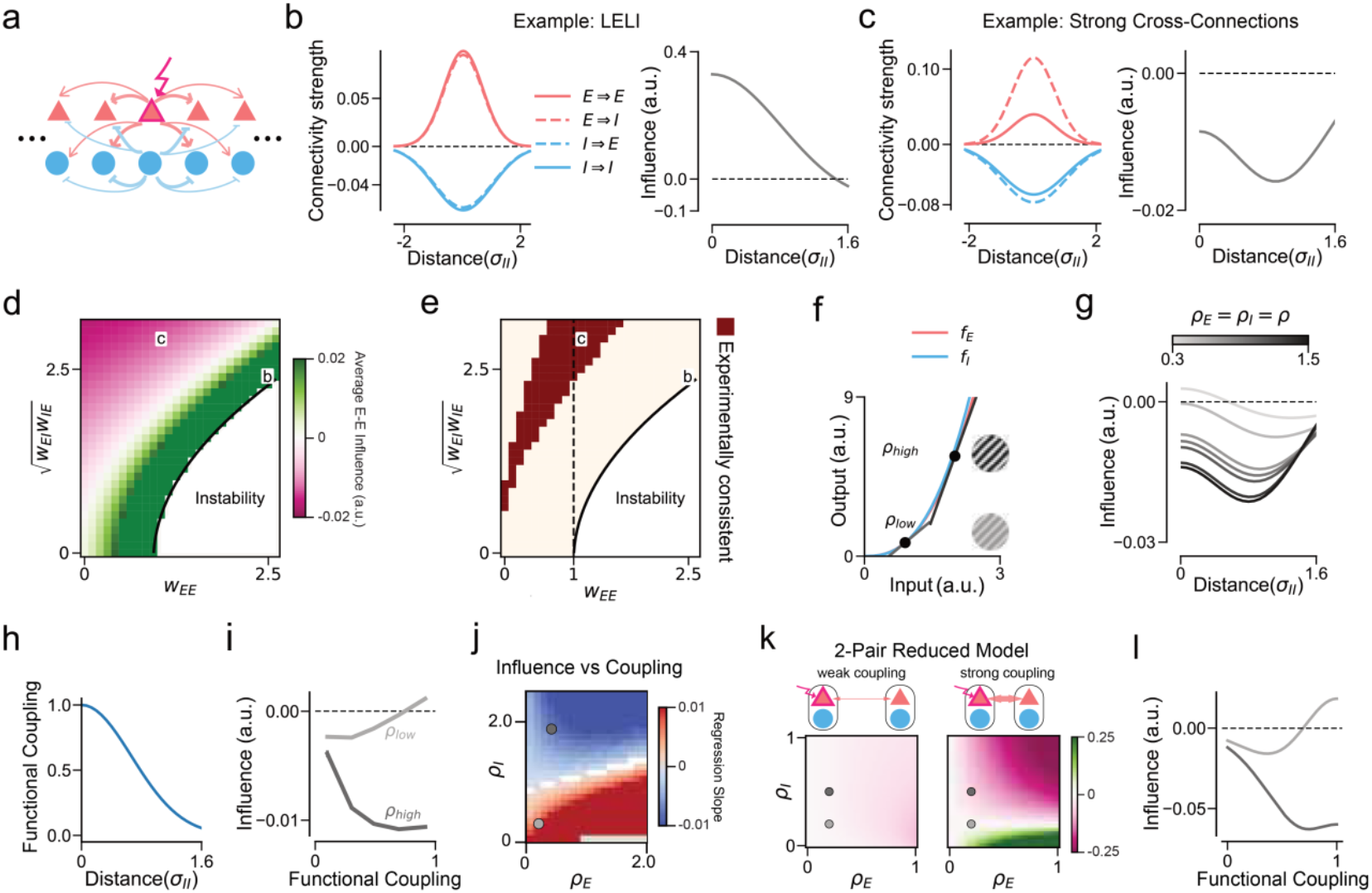
A network model with strong connections between excitatory and inhibitory populations captures experimental excitatory perturbations. (a) Depiction of model architecture: excitatory (E) and inhibitory (I) nonlinear units are connected together defined by Gaussian kernels. We study perturbation of a single excitatory unit at steady-state. (b) Classical LELI connectivity (*left*) leads to large, local facilitatory influences that dissipate over cortical distance (*right*). Here, distance considered is 1.6σ_*I*_, approximately the distance examined in the experimental data. (c) Same as in **b** for stronger cross-population connections (*WEI, W*_*IE*_), compared to within-population (*WEE, W*_*II*_), which produces large, local suppression. (d) Spatially averaged excitatory influence for different connection strengths and a fixed I-I weight (*W*_*II*_ = 2.5). Examples in **b, c** are shown. (e) Area consistent with experimental observations (see Figures 2-3) and ferret visual cortex functional topography. Inhibitory stabilized regime indicated by a black dashed line: regions where *W*_*EE*_ > 1. (f) Network model is linearized at different locations along the power law function set by gain terms (ρ) for E and I populations. (g) Gain modulates spatially-dependent excitatory perturbation suppression. Results shown for model parameters in **c**, choosing identical values for E and I gain. Average influence is negatively correlated with gain (slope = -0.011, p = 0.001). (h) Illustration of functional coupling following normalized E-E connectivity strength γ*E* over distance (see Methods). (i) Excitatory influence depends on functional coupling with nontarget units in a gain-dependent manner. Shown are simulation results for model parameters in **c** at: low gain/contrast (ρ*E* = 0.22, ρ*I* = 0.32) and high gain/contrast (ρ*E* = 0.43, ρ*I* = 1.88). (j) Functional-coupling dependent influence is modulated by E and I gain. Shown is the linear regression slope magnitude and sign calculated on unbinned simulation data for each set of E and I gain values. White regions indicate areas where slope is not significantly different from 0 (Wilcoxon-Mann-Whitney test, p <0.05). (k) *Top*: Reduced model with two E-I neuron pairs with variation in functional coupling between pairs. As in **h**, coupling strength is a function of distance and determines the strength of cross-pair relative to within-pair connections. *Bottom*: Shown are analytically-derived solutions for E-E influence (Eq. 15, see Methods) as a function of gain ρ*E* and ρ*I*. With weak coupling (cross-pair E and I connection strengths reduced by factors γ*E* = 0.012, γ*I* = 0.14, respectively, *left*) and influence is overall negative due to weak direct excitatory connections. With strong coupling (identical cross- and within pair connections, *right*) influence exhibits a gain-dependent switch between facilitation and suppression. (l) As **i**, but for the reduced model in **k** at low gain (ρ*E* = 0.2, ρ*I* = 0.2) and high gain (ρ*E* = 0.2, ρ*I* = 0.5).

Therefore, we next sought to identify the conditions when excitatory units in this network would exhibit broad suppression. Because we are studying influences around a fixed point, we are able to analytically derive how connection weights satisfy an inequality describing when influence is suppressive on average (see Methods and Appendix):

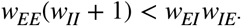

This inequality states that a single excitatory cell exerts average suppression onto excitatory neighbors when cross-population connections (*w*_*IE*_ *w*_*EI*_) are sufficiently large compared to within-population connections (*w*_*II*_ *w*_*EE*_). Under such conditions, we observe suppression consistent with our experimental results (**Fig. 4c**). We then systematically examined how net excitatory influence depends on the strength of within- and cross-population strengths, while keeping *wII* fixed. This revealed three regimes: average suppressive influence, average facilitatory influence, and instability when *wEE* is too large (**Fig. 4d**). These results are consistent across a range of *w*_*II*_ (**Supplementary Fig. 6**). Our experimental results were best matched within a ‘biologically plausible’ zone (**Fig. 4e, Supplementary Fig. 7**) where, in addition to average suppressive influence (1) and stability (2), local cells exhibit suppression (3), and maximum suppression occurs at a non-zero distance (4). We define this zone as the ‘Cross-Dominant’ regime.

Given the ability of a linearized network to capture the suppressive influence of single excitatory units, we wanted to understand how this behavior could change under different stimulus strengths or gain levels. Thus, we linearized the network at different input levels (**Fig. 4f**), resulting in different gain levels of excitatory and inhibitory populations (*ρ*_*E*_, *ρ*_*I*_). In other words, a higher stimulus contrast results in a higher gain^35,36^. Using the network in Figure 4c, we systematically varied gain (with *ρ*_*E*_ = *ρI*), observing enhanced suppression with higher contrast similar to our data (**Fig. 4g**). This change in excitatory influence is also clearly evident when systemically varying connection strengths for different gain values (**Supplementary Fig. 8**).

Finally, we explored how influence depends on functional coupling in our model to examine whether it can explain the switching effect within correlated subnetworks (**Fig. 3c**). We assume the coupling between two excitatory neurons is proportional to the normalized strength of their direct synaptic connections^8,34^, thus exhibiting a distance-dependent Gaussian profile (**Fig. 4h**, see Methods). This allowed us to repeat simulations using the network from Figure 4c while varying excitatory and inhibitory gain (*ρ*_*E*_ and *ρ*_*I*_, respectively). Here, we find that at low gain, influence is positively correlated with functional coupling, switching to a negative relationship at higher gain values (**Fig. 4i-j**). Interestingly, this switch depends primarily on changes in inhibitory gain (*ρ*_*I*_ > *ρ*_*E*_, **Fig. 4j**).

To understand the minimal requirements for this switching behavior, we constructed a reduced model consisting of two excitatory-inhibitory pairs with variable functional coupling between them (**Fig. 4k**, top; see Methods). Closed form solutions of this model (see Appendix) show that for weak coupling, influence is negative, changing little with gain (**Fig. 4k**, *left*). For strong coupling, influence switches depending on gain (**Fig. 4k**, *right*). Specifically, for small *ρ*_*I*_, the strong direct excitatory connection dominates and drives facilitation, whereas for larger *ρ*_*I*_, the disynaptic inhibition across pairs dominates and results in suppression. This dependency on functional coupling is similar to the full model (**Fig. 4l**) and captures several key features in our experimental data (**Fig. 3c**). Note that this model assumes a relation between noise correlation (functional coupling) and excitatory connectivity, but distance dependencies are not required.

As excitatory-driven suppression must originate from inhibitory cells through multisynaptic interactions^7,12,13,37^, we sought to directly test whether inhibitory-to-excitatory (I-E) interactions are consistent with a Cross-Dominant regime and if inhibitory cells strongly suppress excitatory cells. Thus, we performed additional experiments targeting inhibitory cells for two-photon photostimulation using the mDlx promoter^14^, while labeling both excitatory and inhibitory cells with jGCaMP8s (**Fig. 5a, Supplementary Fig. 9**). We measured influences between 44 target inhibitory cells and 967 nontarget putative excitatory cells (without nuclear mRuby2 expression), across 6 FOVs (n = 3 animals). Under both low and high contrast stimuli, I-E influences are significantly more suppressive than excitatory-to-excitatory (E-E), but similar in spatial extent (**Fig. 5b**). In addition, inhibitory suppression onto excitatory cells is invariant to contrast. Modeling inhibitory perturbations in our rate network in a Cross-Dominant regime (**Fig. 4c**) largely captured these observations (**Supplementary Fig. 10**). We also find I-E suppression to increase with noise correlation between cell pairs, except when perturbing an inhibitory target with the preferred visual stimulus at low contrast (**Fig. 5c-d**). During perturbation trials matching a target’s preference, we observed significantly less suppression at low contrast between correlated pairs (r_*noise*_ > 0.5, p < 0.001), potentially revealing the difficulty in suppressing excitatory cells during facilitatory interactions within subnetworks.

**Figure 5.**
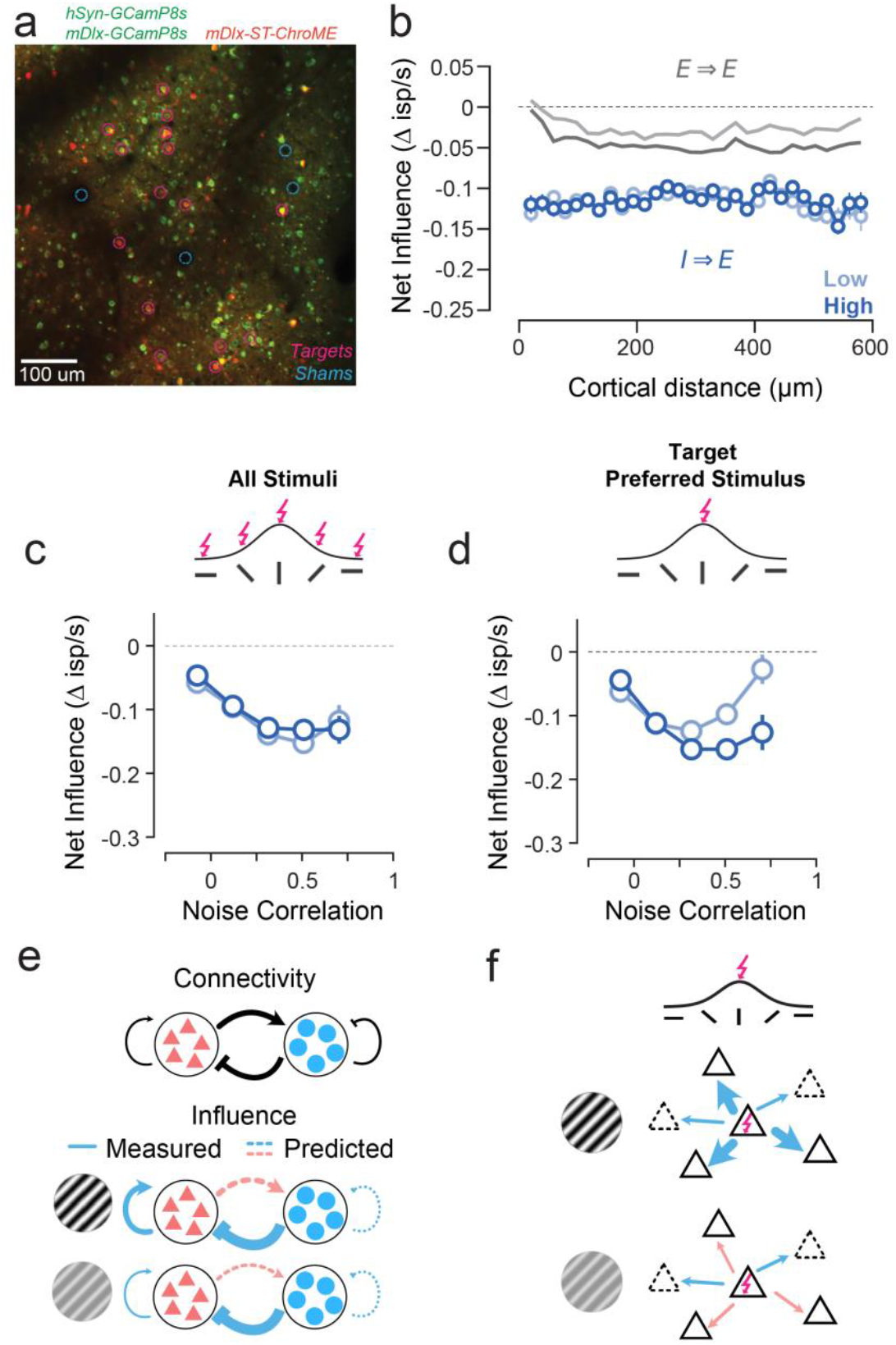
Single inhibitory interneurons provide strong suppression onto excitatory cells. (a) Example field-of-view (FOV) with target (pink) and sham (blue) locations marked. Image is a merge of the average projection (1000 frames) of green and red PMT channels. Cells expressing ST-ChroME (and mRuby2) are inhibitory (driven by mDlx promoter). GCaMP8s expressing cells without mRuby2 expression are putative excitatory neurons. (b) Distant dependent net inhibitory-excitatory (I-E) influence during low and high contrast stimuli (light and dark red, respectively). Distances less than 20 µm were excluded for potential photostimulation artifacts. Each data point represents mean and standard error for that distance bin. Also shown are excitatory-excitatory influences for each contrast (mean values replotted from Fig. 2b). I-E suppression at each contrast is significantly greater than excitatory-to-excitatory (E-E) (p < 0.001 for both). I-E suppression is not significantly different between contrasts (p = 0.16). (c) Relationship between local net influence and noise correlations for low (light red) and high (dark red) contrast. Data points are mean and standard error across all photostimulation trials. Spearman’s *r*_*high contrast*_ = - 0.16, n = 16,213 comparisons, p < 0.001. Spearman’s *r*_*low contrast*_ = -0.21, n = 16,307 comparisons, p < 0.001. (d) Same as in **c** for photostimulation trials aligned with the target cell’s preferred orientation (< 30 deg). Spearman’s *r*_*high contrast*_ = -0.17, n = 2,450 comparisons, p = < 0.001. Spearman’s *r*_*low contrast*_ = -0.044, n = 2,491 comparisons, p = 0.12. (e) Cross-Dominant connectivity has stronger connections between excitatory and inhibitory populations. Strong I-E connectivity provides a circuit mechanism underlying spatially-broad perturbation-driven suppression. Recurrents activated during low contrast stimulation (e.g. low gain) are less suppressive than during high contrast stimulation. (f) Functionally-coupled excitatory subnetworks exhibit feature competition at high contrast and feature amplification at low contrast during strong feed-forward activation.

## Discussion

Altogether, our study suggests that the ferret visual cortex operates in a Cross-Dominant regime, where excitatory influence primarily exhibits spatially-broad feature-competition (suppression) and gain changes can switch influence between functionally-coupled excitatory cells (**Fig. 5e-f**). Analysis of a rate-based recurrent neural network consisting of excitatory and inhibitory nonlinear units show many of our experimental results from single excitatory and inhibitory cell perturbations can be explained by excitatory-inhibitory connections being considerably stronger than connections between excitatory cells. Notably, this Cross-Dominant regime is different from a classic LELI structure, in which strong excitatory connections within columns produce modular activity patterns^27–30^. Although, there is evidence that the latter more accurately describes early cortex^27,30^, when modular activity is most pronounced^38^. Our results are also consistent with a Stabilized Supralinear Network^5^: when excitatory-to-excitatory connections are sufficiently strong, at high gain (contrast), the network operates in an inhibition stabilized regime^2,5^. In addition, we show that the gain-dependent switching behavior can be understood within a reduced model of two pairs of E/I units.

We note that our model does not perfectly capture our empirical data, which could be due to missing visual circuit details. Our models do not include long-range connectivity^39^ or local heterogeneity, and we did not consider the rich feature space encoded by V1. In addition, as genetic tools are currently being developed to target inhibitory cell subtypes in nonmurine species, we were not able to perturb their activity experimentally and thus chose to not consider subtypes in our network models. Nonetheless, this work emphasizes the importance of connections between cell types and functional subnetworks within functional, modular circuits.

Cross-Dominant cortical connectivity may be a universal property of the mammalian neocortex. Slice physiology in human and mouse cortex shows far greater synaptic connectivity between excitatory and inhibitory cells compared to excitatory alone^40^. Such a connectivity motif is consistent with a like-to-like organization between excitatory and inhibitory cells^41^. However, our findings do deviate from experimental data and modeling from mouse V1^6,7,11–13^. First, we do not observe strong amplification directly nearby perturbed excitatory cells, but rather broad feature-competition that is often greater in magnitude than amplification (**Fig. 2**). Comparing spatial influence profiles, mouse V1 appears more consistent with LELI-like influence, although generating such influence does not require LELI connectivity^13^. Thus, functional microclusters in mouse V1^8^ are unlikely to be homologous with functional columns in ferret. Second, we observe stimulus-dependent modulation, not yet observed in mouse V1, but predicted by mathematical analysis of models of single-cell perturbations^13^ and *in silico* ensemble perturbations^11^. Interestingly, the strong I-E influences that we predicted and observed are consistent with models of mouse V1 showing that functionally specific inhibitory connectivity underlies feature-specific suppression^12^. Third, we observed stronger influences between correlated pairs of cells (**Fig. 3**), more than pairs with only similar tuning (**Supplementary Fig. 5**). In contrast, competition in mouse V1 was observed between cells with similar tuning, not between those with strong functional coupling (which where dominated by amplification). This suggests that in mouse V1, separate subnetworks could be responsible competition and amplification, while we find that a single network flexibly switches in a stimulus-dependent manner.

By linking functional, modular organization with recurrent connectivity motifs, our work bridges experimental and theoretical frameworks, offering a mechanistic foundation for how sensory signals can, depending on context, dynamically reshape cortical computations. At the same time, our findings expose a critical gap: the structure and specificity of excitatory– inhibitory connectivity remain poorly understood, yet are likely central to flexible computations of cortical circuits.

## Acknowledgements

We thank Tammi Coleman for helping to support animal surgeries and experiments, the animal support staff at University of Colorado at Anschutz Medical Campus, Greg Bond for helping to establish and maintain our image processing pipeline, the GENIE project for access to GCaMP8, Dr. Hillel Adesnik for sharing ChroME constructs for preliminary experiments and adapted constructs. We thank Drs. David Fitzpatrick and Alon Poleg-Polsky for helpful discussion and comments.

## Funding

National Institutes of Healthy grant R00EY031137 (BS)

National Institutes of Healthy grant R21EY034992 (BS)

Whitehall Foundation Award (BS)

Alfred P. Sloan Foundation Fellowship Award (BS)

Bundesministerium für Bildung und Forschung grant 01GQ2002 (MK,DK,LB)

Deutsche Forschungsgemeinschaft Research Unit FOR 5368 ARENA (MK,DK)

## Author contributions

Conceptualization: BS Methodology: BS, DK, MK Investigation: JB, BS, DK, LB, MK Visualization: BS, DK, LB Funding acquisition: BS, MK Project administration: BS, MK Supervision: BS, MK Writing — origin draft: BS, DK, LB, MK Writing — review & editing: BS, DK, LB, MK

## Data, code, and materials availability

Data and code are available from the corresponding author upon reasonable request.

## Supplementary Materials

Methods

Appendix

Figs. S1 to S10

## 1. Methods

All procedures were performed according to NIH guidelines and approved by the Institutional Animal Care and Use Committee at the University of Pennsylvania and University of Colorado Anschutz Medical Campus.

### 1.1. Viral Injections

Female and male ferrets (n = 12) aged P14-21 (Marshall Farms) were anesthetized with isoflurane (delivered in O2 with a Somnosuite). Bupivacaine was administered SQ. Animals were maintained at an internal temperature of 37^*o*^ Celsius. Under sterile surgical conditions, a small craniotomy (0.8 mm diameter) was made over the primary visual cortex (5-8 mm lateral and 2-4 mm anterior to lambda). One of two mixtures of AAV was injected (125 - 202.5 nL) through beveled glass micropipettes (10-15 µm outer diameter) at 600, 400, and 200 µm below the pia. The first mixture (to target excitatory cells) contained: AAV1/2.hSyn.Cre, AAV1/2.hSyn.GCaMP8s^42^, and AAV9.CAG.FLEX.ChroME-ST.P2A.H2B.mRuby2^19^.The second mixture (to target inhibitory and excitatory cells) contained: AAV1/2.hSyn.GCaMP8s, AAV1/2.mDlx.GCaMP8s^14^, AAV1/2.mDlx.ChroME-ST.P2A.H2B.mRuby2. Finally, the craniotomy was filled with sterile agarose (Type IIIa, Sigma-Aldrich) and the incision site was sutured or glued (Vet Bond).

### 1.2. Cranial Window

After 4-8 weeks of expression, ferrets were anesthetized with 10-15 mg/kg ketamine and isoflurane. Atropine and bupivacaine were administered, animals were placed on a feedback-controlled heating pad to maintain an internal temperature of 37^*o*^ Celsius, and intubated to be artificially respirated. Isoflurane was delivered throughout the surgical procedure to maintain a surgical plane of anesthesia. An intravenous cannula was placed to deliver fluids. Tidal CO2, external temperature, and internal temperature were continuously monitored. The scalp was retracted and a custom titanium headplate adhered to the skull with dental cement. A craniotomy was performed and the dura retracted to reveal the cortex. One piece of custom coverglass (4.5 mm diameter, 0.7 mm thickness, Warner Instruments) adhered to a custom insert using optical adhesive (71, Norland Products) was placed onto the brain to dampen biological motion during imaging. A 1:1 mixture of tropicamide ophthalmic solution (Akorn) and phenylephrine hydrochloride ophthalmic solution (Akorn) was applied to both eyes to dilate the pupils and retract the nictitating membranes. Contact lenses were inserted to protect the eyes. Upon completion of the surgical procedure, isoflurane was gradually reduced and pancuronium (2 mg/kg/hr) was delivered IV.

### 1.3. Visual Stimuli

Visual stimuli were generated using Psychopy^43^. The monitor was placed 25 cm from the animal. Receptive field locations for each field of view (FOV) were hand mapped. For each FOV we presented gratings (square and sinusoidal) drifting at 8-16 directions (0 – 315^*o*^) at two contrasts (low: 8-16%, high: 64-100%). Gratings were all presented at 0.08 cycles per degrees and 4 Hz temporal frequency presented to the contralateral eye. Stimuli were presented for 1 second.

### 1.4. Two-photon Imaging and Photostimulation

Two-photon imaging was performed on a Ultra 2P Plus imaging system (Bruker) running PrairieView with a separate photostimulation module and the Neuralight SLM-based module. Imaging was done at 925 nm with Coherent Ultra-2 tunable laser and photostimulation was done with a Coherent Fidelity-2 laser (1070 nm). For imaging, power after the objective was limited to *<* 40 mW. Single-cell photostimulation was optimized for each FOV and ranged 15-60 mW average power per cell (**Extended Data Fig. 1**). FOVs were chosen based on healthy expression of GCaMP8s, the presence of photo-activated cells, and a large number of photostimulation targets to choose from. Images were collected at 30 Hz using bidirectional scanning with 512×512 pixel resolution. Photostimulation was performed with spiral scanning following details in Chettih and Harvey (2018). Two-photon photostimulation was driven following visual stimulus presentation with a ∼ 120 ms delay and consisted of bursts for 250-500 ms. Each burst consisted of 12 galvo-galvo spirals over a duration of 27 ms, with an inter-pulse interval of 23 ms. Small, natural movements of cells due to heart rate and breathing meant that spiral scans did not exactly occur over the same location of a target repeatedly. Photostimulation spots (‘Mark Points’) consisted of targeted cells expressing mRuby2 in the nucleus and sham locations (e.g. blood vessels, neuropil, cortical regions without fluorescence expression at or above or below the focal plane). For each FOV, 1-4 sham locations were chosen, depending on the number of targets.

### 1.5. Two-Photon Imaging Processing

Imaging data were excluded from analysis if motion along the z-axis was detected. In-plane motion was corrected using rigid registration and a piecewise non-rigid motion correction algorithm^44^ run in Python. Following registration, data were denoised using DeepInterporation to increase signal-to-noise^45^. Cellular ROIs were drawn in ImageJ using Cell Magic Wand^14^. Mean pixel values for ROIs were computed over the imaging time series, Δ*F/F*_*o*_ was computed by computing *F*_*o*_ with time-averaged median or percentile filter (10th percentile), and then underlying spike rates were inferred using CASCADE^26^. Activity traces were synchronized to stimulus triggers sent from Psychopy and collected by Bruker’s recording acquisition system. Peak responses (inferred spikes/sec) to visual stimuli were computed each trial to capture the full stimulus-driven modulation. Cells used for subsequent analyses were required to have peak responses whose distribution exhibited greater skewness than the average skewness of sham ROIs^41^.

### 1.6. Generalized Linear Model (GLM) to Quantify Influence

For each neuron, we fit a Poisson GLM to predict its trial-by-trial responses. The model includes four types of predictors: (1) visual stimulus identity, (2) optogenetic perturbation to target neuron (3) perturbation to sham locations, and (4) global activity (cell population average trial-by-trial activity).

The GLM is formulated as:

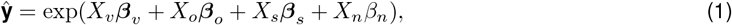

where **ŷ** (*N*_*trials*_ × 1) is a vector denoting the predicted response of that neuron in all trials. *X*_*v*_, *X*_*o*_, *X*_*s*_ are binary design matrices (*N*_*trials*_ × *N*_*stimuli*_), denoting the presence of visual grating, optogenetic target and sham stimuli, respectively. *X*_*n*_ (*N*_*trials*_ × 1) is a continuous regressor capturing the global activity level, defined as the average activity over all neurons within the FOV per trial. We fit the model weights 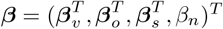, by minimising the squared error of firing rate prediction using coordinate descent, with elastic net regularization

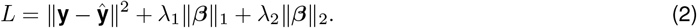

Five-fold cross-validation (80% training, 20% test split) was used to determine the optimal regularization parameters *λ* _1_, *λ* _2_ for each neuron. Model fitting was performed in Python (version 3.12.4) and the statsmodels package (version 0.14.2).

We assumed that perturbing sham locations does not alter the firing of non-target cells. Therefore, sham weights were used to estimate and correct bias in the opto-target weights. For each target neuron *t*, we generated 20 bootstrap resamples of the data and computed the distributions of fitted target weights (*b*_*t*_, *t*-th component of ***τ***_*o*_) and sham weights (*b*_*s*_). We then subtracted the mean sham weight 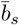 over all bootstrap samples and sham locations from inferred target weights *b*_*t*_ and average over bootstrap samples to obtain bias-corrected estimate 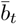. To assess significance of target-nontarget weights, we computed d-prime

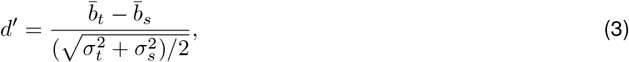

where *σ* _*t*_, *σ* _*s*_ are standard deviations of the target and sham distribution, respectively. We considered target weights significant if |*d* ^′^ | *>* 1.

Significant opto-target weights were converted into an “influence” measure expressed as the change in inferred spikes/sec. For each nontarget neuron, we compared the GLM prediction of the full model and the prediction with opto-target weight set to 0. Influence of target cell *t* was then computed as difference between these two predictions, averaged over trials

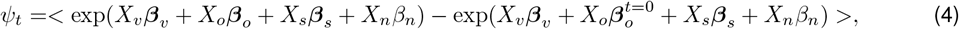

where *<>* denotes averaging over trials, and 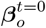 is the fitted weight vector with target *t* set to 0. To obtain the stimulus dependent influence, we fit the GLM only on trials when only the stimulus is shown. To quantify the ability of a GLM in capturing a nontarget cell’s trial-by-trial activity, we computed the pseudo *R*^2^(deviance explained) per cell, defined as

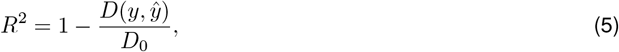

where

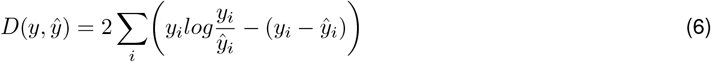

is the deviance of the GLM, and

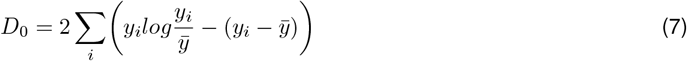

is the deviance of the null model that only captures the mean activity of a cell over trials.

### 1.7. Analysis and Statistics

Targets were only used for subsequent analyses if they had robust photostimulation activation (**Extended Data Fig. 1**). Influences between target and nontarget cells were only used if significantly different than the sham locations in the same FOV (see above). Otherwise, measured influences were ignored. Out of 181 excitatory cell targets, 133 passed criteria and were used for subsequent analysis. Across all FOVs, we collected 161.0 ± 48.5 (mean ± s.d.) photostimulation trials per target and across all targets, 10.25 ± 4.8 (mean ± s.d.) trials per visual stimulus class. Out of 59 inhibitory cell targets, 44 passed criteria and were used for subsequent analysis. Across all FOVs, we collected 175.4 ± 25.3 (mean ± s.d.) photostimulation trials per target and across all targets, 11.0 ± 3.4 (mean ± s.d.) trials per visual stimulus class.

Trial-by-trial correlations and GLM tuning similarity were calculated by the Pearson correlation coefficient of peak inferred spike rate across trials between all cells. Noise correlations were calculated by measuring the Pearson correlation coefficient between stimulus-specific mean-subtracted residuals. Direction/orientation preference for each cell was estimated by fitting a double Gaussian to each cell’s mean visual responses during sham photo-stimulation trials^46^. Statistical tests were performed in Matlab and Python. We used nonparametric statistical tests when data was not normally distributed. For comparisons between two distributions, a rank-sum test was used unless otherwise stated.

### 1.8. Full Network Model

A comprehensive description of the network model used to derive the results in **Fig. 4** and related **Extended Data Figures** are provided in the **Appendix**. In the following, we present the main steps that led to our results.

To characterise how properties of recurrent excitation and inhibition, under different visual stimulus conditions, shape the influence of single-cell optogenetic stimulations on surrounding neurons in a network, we consider a continuous one-dimensional firing-rate model of recurrently connected excitatory (E) and inhibitory (I) populations. In the experiments, optogenetic stimulation induces only small activity changes in surrounding cells; we therefore model its effect as a weak perturbation of the steady-state response to the visual input. Accordingly, we analyze the network dynamics linearized around this fixed point, which reads

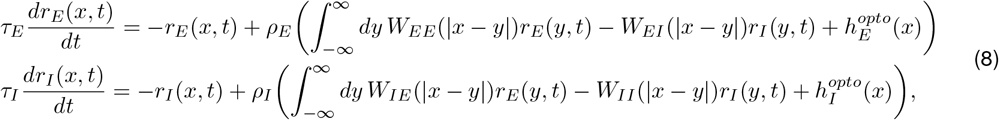

where *r*_*E*_(*x, t*) and *r*_*I*_ (*x, t*) denote the excitatory and inhibitory firing rates, respectively, 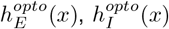 the external inputs, and *τ*_*E*_, *τ*_*I*_ the time constants. For simplicity, we set *τ*_*E*_ = *τ*_*I*_ = 1. *ρ*_*E*_ and *ρ*_*I*_ are respectively the excitatory and inhibitory gain, the change in output rate per change in input, i.e., the I/O function’s slope. We first consider *ρ* _*E*_ = *ρ* _*I*_ = 1. The synaptic weights *W*_*KL*_(|*x* − *y*|) from location *y* in population *L* to location *x* in population *K*, where *K, L* ∈ [*E, I*], are modeled by Gaussians,

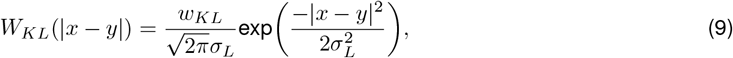

where *w*_*KL*_ is the connectivity strength and *σ* _*L*_ controls the spatial range of the connections, assumed to depend only on the presynaptic neuron and we assume *σ* _*E*_ *< σ* _*I*_. The optogenetic stimulation to a single excitatory neuron is modeled as a Dirac delta centered at *x*_0_ = 0 and with a amplitude *h*_*E*_ *>* 0

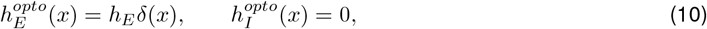

with the inhibitory input set to zero, and vice versa when modeling the stimulation of inhibitory neurons (**Extended Data Fig. 10**).

At steady state, Eq. (8) reads in more compact form

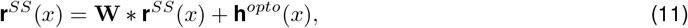

where 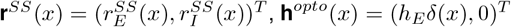, **W** is a 2 × 2 matrix with elements *W*_*KL*_(|*x* − *y*|), and ∗ denotes convolution. Solving Eq. (11) in the Fourier domain and isolating the part that depends on the recurrent connectivity (i.e. subtracting out the part that depends on the input only), the E-E (E to E) influence yields

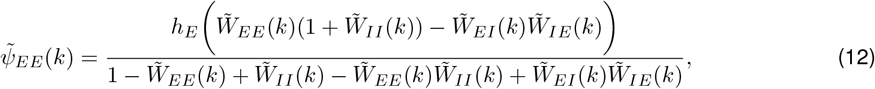

where 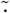 denotes Fourier transform. The E-E Influence in real space (**Fig. 4b-c**) is obtained by taking the inverse Fourier transform of Eq. (12). We determine the net E-E influence 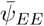 by evaluating Eq. (12) at *k* = 0. Requiring the solution to be stable, the denominator must be positive, and therefore net suppression requires

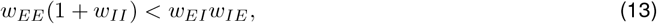

defining the Cross-Dominant regime. The ‘biologically plausible’ zone (**Fig. 4e)** is the parameter regime defined by two further conditions, namely that E-E influence is locally suppressive,

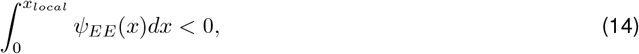

where we used *x*_*local*_ = 0.4*σ*_*I*_ and that suppressive influence is strongest at a distance larger than zero from the stimulated neuron,

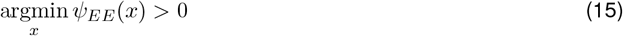

(see also **Extended Data Fig.7**).

To study the effect of visual contrast on influence, we assume a power law nonlinearity following previous studies^2,36^. For such nonlinearity the excitatory gain *ρ*_*E*_ and inhibitory gain *ρ*_*I*_ increase with visual drive, which is assumed to increase with contrast (illustrated in **Fig. 4f**). With excitatory and inhibitory gain, the influence (Eq. 12) reads

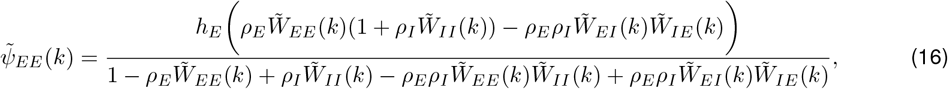

and the inverse Fourier transform of this expression is plotted in **Fig. 4g** for various values of gain *ρ*_*I*_ = *ρ*_*E*_ = *ρ*. The condition for suppressive net influence now yields

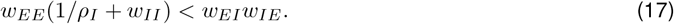

showing that the influence depends on the inhibitory gain only, and tends to be suppressive for strong gain, while it can become positive for small gain.

To model the gain-dependent switch in functionally coupled subnetworks observed in the experiments (**Fig. 3c**), we assume functional coupling is proportional to the strength of direct excitatory connections, normalized to a maximal value of 1. Thus, functional coupling is equal to the relative strength of E-E connections,

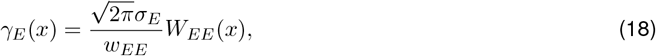

with 1= *γ*_*E*_(0) ≥ *γ*_*E*_(*x*) ≥ 0. It is depicted in **Fig. 4h** and used to depict the relationship between E-E influence and functional coupling for different combinations of gain values *ρ*_*E*_ and *ρ*_*I*_ in **Fig. 4j**.

### 1.9. Reduced Model

We constructed a reduced model of two coupled pairs of neurons, each pair consisting of an E and I neuron. First, we consider all connections of the same type (from population *K* to *L*, with *K* and *L* ϵ [*E, I*]) to be identical, forming a minimal model of a strongly coupled subnetwork. Stimulating one of the two excitatory neurons, the influence on the other one is

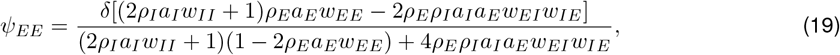

where *δ* is the input amplitude, the parameters 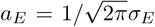 and 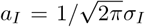 are introduced to facilitate the comparison with the full model. This influence is suppressive if

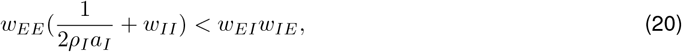

an expression similar to Eq. (17). We then vary the coupling strength between the two pairs and compute influence as a function of the gains *ρ*_*E*_ and *ρ*_*I*_, assuming the cross-connections between pairs are symmetric. As in the full model, the excitatory cross-connections are reduced by a factor 0 ≤ *γ*_*E*_ *≤* 1 equal to the coupling strength. We assume that inhibitory connections decrease along with the excitatory connections, and that their ratio is the same as in the full model. Thus, we have

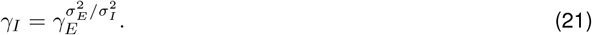

Note that, sinceσ_*E*_ *< σ*_*I*_, the decrease of inhibitory connections is weaker than that of the excitatory connections. The influence as a function of gain and functional coupling (**Fig. 4k-l**) is given by

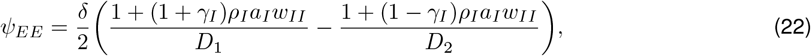

where *D*_1_ and *D*_2_ are given in the **Appendix**.

## Appendix to “Single-cell perturbations reveal adaptive, stimulus-dependent recurrence in visual cortex”: Extended model description

## 1. Network model of E-E influences

### 1.1. Linear network model

To characterize how recurrent excitation and inhibition shape influences in ferret visual cortex, we consider a continuous firing-rate model of recurrently connected excitatory (E) and inhibitory (I) populations. Influences are the neural responses of neurons in the network to single-cell optogenetic activation. In our experiments, optogenetic activation was performed during visual stimulation with moving gratings at different contrasts. Assuming that such single-cell input induces only a weak perturbation of the visually evoked steady-state response in surrounding neurons, we analyze the network dynamics linearized around this steady state. Using a compact vector formulation, the linearized dynamics are written as

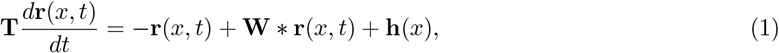

where 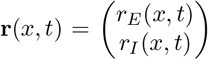, denotes the firing rates of E and I populations, respectively, 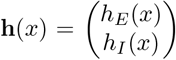 represents the external input (i.e. the optogenetic activation, assumed constant in time) and 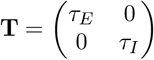 is the matrix of time constants. The symbol ∗ denotes spatial convolution

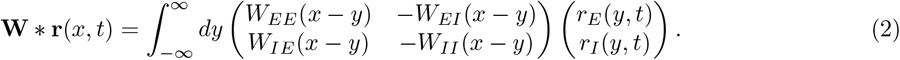

We assume that the connectivity kernels *W*_*KL*_(*x y*) are ≥ 0 and depend only on the distance between the spatial locations *x* and *y* for all *K, L* ∈{*E, I*} and not on the specific location in the network.

Each kernel is normalized such that it spatial integral

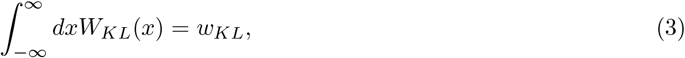

where *w*_*KL*_ denotes the total effective connectivity strength from population *L* to population *K*.

### 1.2. Gaussian connectivity

Concretely, in Fig. 4b-g, we used Gaussian connectivity kernels

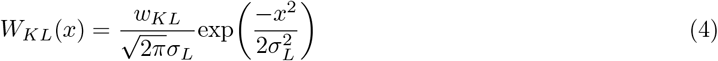

with *σ*_*I*_ *> σ*_*E*_ to model how influences depend on spatial distance and visual contrast. Such Gaussian kernels approximate the spatial decay of synaptic connections in ferret visual cortex. Evidence for spatially extended inhibitory connections comes from [7]. Choosing the weights *w*_*KL*_ appropriately, this connectivity motif can give rise to local excitation and lateral inhibition [8] [2](LELI; sometimes also referred to as a “Mexican hat” profile). Previous studies have shown that rate models of type Eq. (1) with such connectivity can capture the modular patterns of spontaneous and visually evoked activity in the developing ferret visual cortex [1], [2], [9]. For all plots in Fig. 4b-g we used *τ*_*I*_ = 1.5*τ*_*E*_. Note, however, that most of the results presented below do not assume a specific functional form of the connectivities *W*_*KL*_(*x*) other than condition Eq. (3).

### 1.3. Steady-state solution

To obtain the steady-state response, we apply the Fourier transform to Eq. (1) and use the convolution theorem

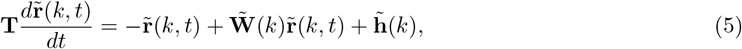

where 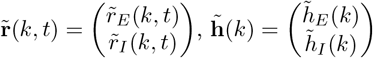 and 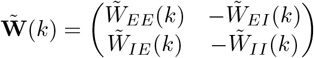, and

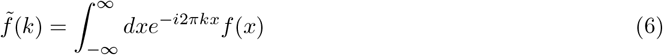

defines the Fourier transform of function *f* (*x*). For simplicity, we assume identical time constants for both populations, in which case we can set 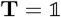.

At steady state, defined by 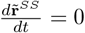, the solution to Eq. (5) is given by

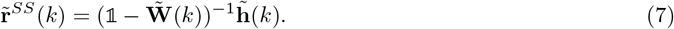

Rewriting Eq. (7) allows us to separate the part that involves the recurrent interactions from the part that only depends on the stimulation:

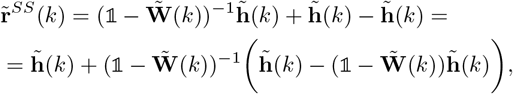

and thus

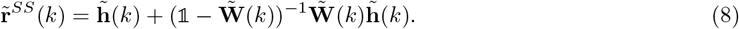

The steady-state response to the optogenetic activation, Eq. (8), thus consists of a feedforward input component plus a recurrently amplified component. This separation allows us to isolate the pure recurrent influence in Section 1.4.

### 1.4. Stability condition

For Eq. (5) to be stable, the real part of the maximum eigenvalue of the Jacobian 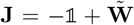 must be negative. This condition requires Det(**J**) *>* 0 and Tr(**J**) *<* 0, which yields

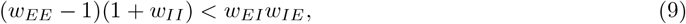

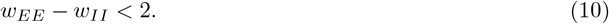

This stability condition imposes constraints on the connectivity strength parameters *w*_*KL*_.

### 1.5. E-E influence

This section describes how E–E influence is modeled. In the experiments, we deliver optogenetic stimulation to a single excitatory neuron. We model this stimulation as an external input proportional to a Dirac delta centered at *x*_0_, the location of the stimulated neuron. Without loss of generality, we set *x*_0_ = 0. The Fourier transform of this input is

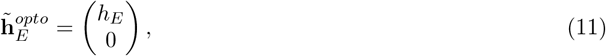

where *h*_*E*_ *>* 0 denotes the amplitude of the optogenetic input in Fourier space, and the inhibitory component is zero, focusing on the case, where only the excitatory neuron is stimulated. Given this input, we define “influence” as the steady-state response

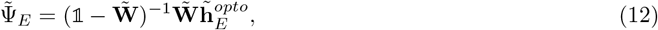

to the stimulation 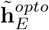, restricted to the part that depends on the recurrent interactions (see Eq. (8)). The E to E (E-E) influence in the Fourier space is then given by the first component of 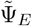, and yields

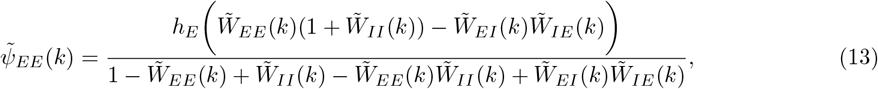

where we used

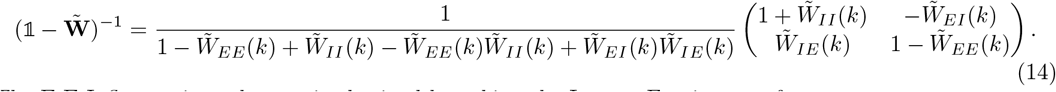

The E-E Influence in real space is obtained by taking the Inverse Fourier transform

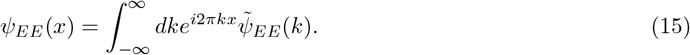

Eq. (15) describes the E-E influence as a function of spatial distance *x* allowing for a direct comparison between the experimental observations and the model.

### 1.6. Net E-E influence

We compute the net E-E influence 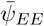 by averaging *τ*_*EE*_(*x*) over all network positions

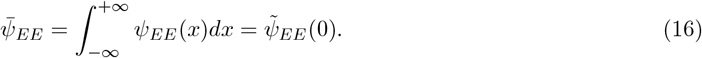

Owing to the normalization condition in Eq. (3), we have

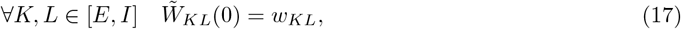

and using Eq. (13) the net E-E influence yields

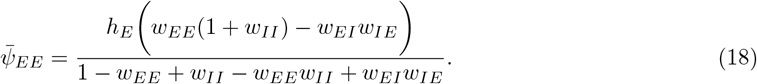

This equation determines the conditions under which the net E–E influence is suppressive, as observed in our experiments (Fig. 2). Because of the stability requirement of the network in Eq. (9), the denominator of Eq. (13) is strictly positive. Consequently, the sign of the net E–E influence is determined solely by the numerator. Since *h*_*E*_ is positive, the numerator is negative if

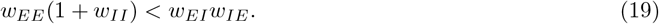

Eq. (19) establishes the condition on the connectivity strengths for a net E–E influence that is suppressive.

### 2. Synaptic expansion

To better understand the spatial dependence of E-E influence, we express the influence as a function of space using an expansion in increasing orders of synaptic interactions. Following [3] and [4], we express the steady-state solution **r**^*SS*^(*x*) in real space as the inverse Fourier transform of Eq. (7)

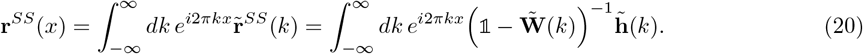

The excitatory component of Eq. (20) is

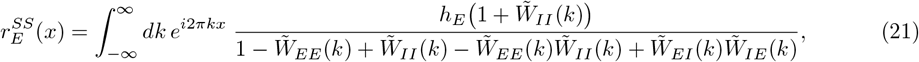

and defining for convenience

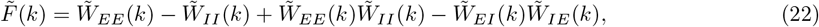

we can write Eq. (21) as

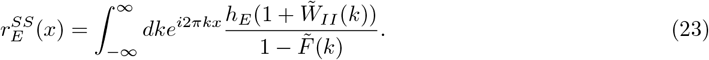

If the Neumann series convergence criteria 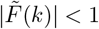 is satisfied, we can expand the denominator as

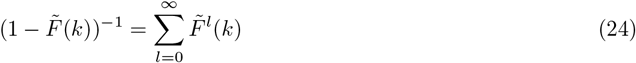

Substituting Eq. (24) into Eq. (23) we obtain

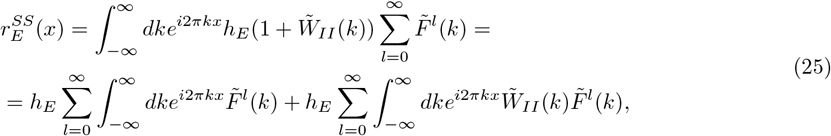

Assuming *h*_*E*_ = 1, we expand Eq. (25) and express it as a sum of mono-synaptic, di-synaptic, tri-synaptic, and higher order terms, expressing the influence as different orders of synaptic interactions

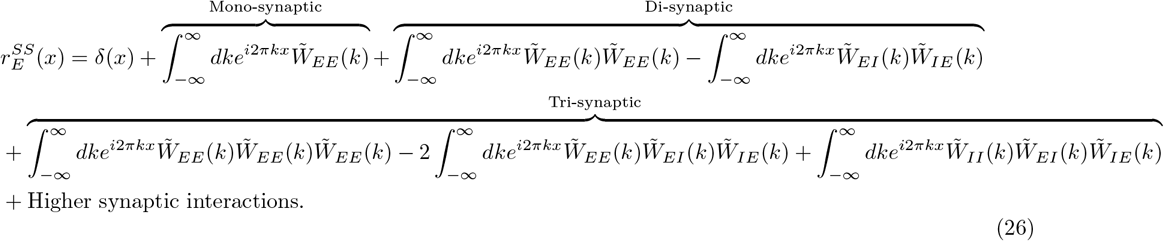

This expansion provides a decomposition of the E-E influence into contributions from pathways of increasing synaptic orders. By expressing the influence as a sum of mono-, di-, and higher-order pathways, the formulation reveals how excitation and inhibition shape the spatial profile of influence. Although this framework can only be applied to a limited parameter range due to the convergence criteria of the Neumann series, it offers an intuitive and analytically tractable way to understand the spatial profile of E-E influence.

## 3. Parameter regime consistent with observed E-E influence

The spatial E-E influence profile revealed by our experiments exhibits further characteristics, in addition to being suppressive on average. In the main text (Fig. 4e) we define four criteria for the influence of our model to be “biologically plausible”, and show the parameter regime in which all criteria are satisfied. The first two criteria – network stability and suppressive net influence – are defined by Eqs. (9), (10) and (19). In the following, we define the other two criteria.

### 3.1. E-E Influence is locally suppressive

As shown in the main text (Fig. 2), the E-E influence is mostly suppressive also in the vicinity of the stimulated neuron. We quantify this by considering the influence up to a distance *x*_*local*_ corresponding to a region within 150*µ*m of the stimulated neuron. The criteria that E-E influence is locally suppressive is defined by

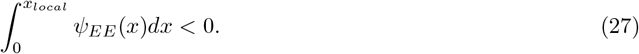

Given the Gaussian connectivity used in our model, 150*µ*m correspond approximately to *x*_*local*_ = 0.4*σ*_*I*_. As we show in the Extended Data Fig. 7, among the parameters that show average suppression, only a subset also exhibits local suppression. Condition Eq. (27) thus imposes further constraints on the experimentally consistent parameters space.

### 3.2. Maximum suppression occurs at a non-zero distance

In the experiments we observe that the suppressive influence is strongest not close to the stimulated neuron, but some distance away. This condition requires that *τ*_*EE*_(*x*) has its minimum at some *x*_*min*_ *>* 0

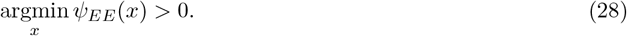

This condition further restricts the parameter space, yielding a reduced subset that is consistent with the experimental observations, as shown in the Extended Data Fig. 7.

## 4. Modeling contrast-dependent gain changes

To describe how changes in contrast affect the spatial influence profiles (Fig. 4f, and g), we consider a nonlinear firing-rate model with a power law nonlinearity following [5] and [6]. Using the same notations as in Eq. (1) this model is defined as

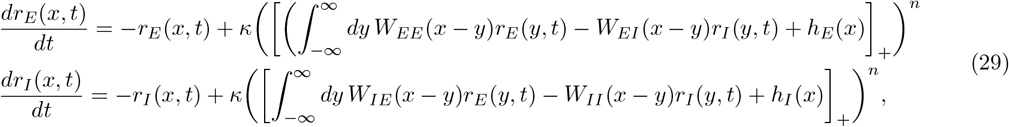

where *σ* is a constant, *n >* 1 is an integer and [*x*]_+_ = max *x*, 0. The inputs *h*_*τ*_(*x*), with *α ϵ* [*E, I*] as before, now reflect both the visual stimulation and the optogenetic activation. Assuming *n* is an integer simplifies the linearization of Eq. (29), revealing the input-dependence of the gain. For a visual input **h**^*V*^ (assumed to be constant in space, for simplicity) the fixed point of Eq. (29) can be obtained through

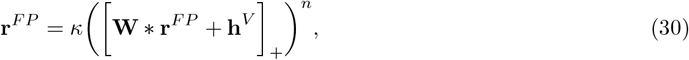

where we used the compact formulation introduced in Eq. (1). We linearize Eq. (29) to obtain the response *δr*_*α*_(*x, t*) to an optogenetic stimulation *δh*_*α*_(*x*) around this fixed point. This is achieved by inserting

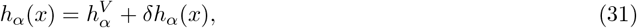

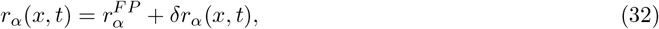

into Eq. (29), where *δh*_*τ*_(*x*) is the optogenetic perturbation defined in Section 1.5. Neglecting quadratic and higher order terms, this yields

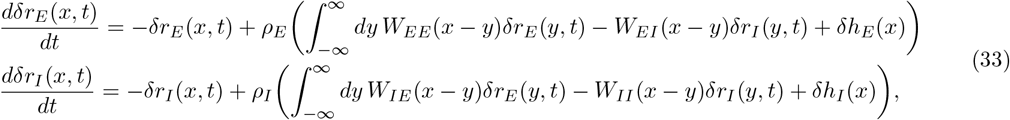

where

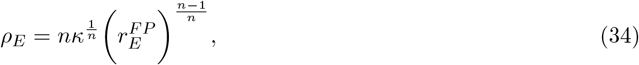

is the excitatory gain and

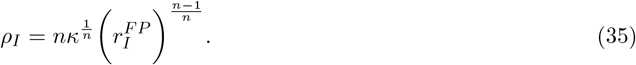

is the inhibitory gain, which both depend on the strength (contrast) of the visual stimulus 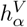 through the response 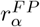. To make the notation simpler we adopt the following change of variables,

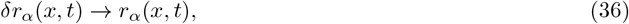

and

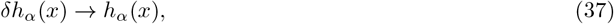

for *α*∈ [*E, I*]. Eq. (33) can be written in the same compact matrix form as Eq. (1) and solved at steady state in the Fourier domain as in Section 1.5. In doing so, we obtain an expression analogous to Eq. (13), but now depending also on the excitatory and inhibitory gains

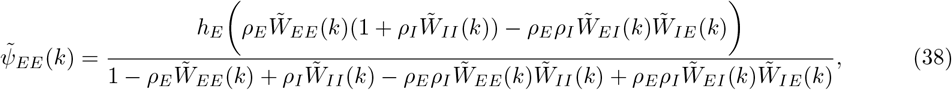

As before, the E to E Influence in real space is obtained by taking the Inverse Fourier transform of this expression.

### 4.1. Net E-E influence depending on gain

The net E-E influence for different gain levels is obtained by evaluating Eq. (38) at *k* = 0

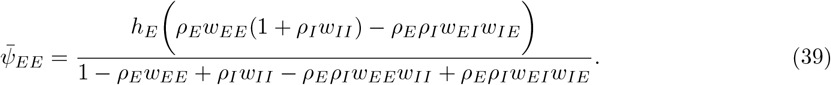

With the simplifying assumption *ρ* = *ρ*_*E*_ = *ρ*_*I*_ used in Fig. 4g, the condition for net suppression is

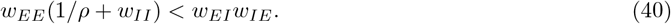

In the more general case, *ρ*_*E* ≠_ *ρ*_*I*_, the condition for net suppression is

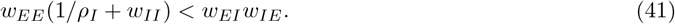

Note that this condition only depends on *τ*_*I*_, but not on *τ*_*E*_.

## 5. Functionally coupled networks

In this section, we examine the relationship between functional coupling and influence (Fig. 4h-l), in order to better understand the gain-dependent switch in functionally coupled subnetworks observed in the experiments (Fig. 3c). In the experiments, noise correlation was used as a proxy for functional coupling. In our model we assume functional coupling is proportional to the strength of direct excitatory connections.

### 5.1. Full model

First, we study how influence depends on functional coupling using the full model introduced in Section 1.1. This is related to Fig. 4h, and i. In our model, the strength of E-E connections *W*_*EE*_(*x*) falls off as a function of distance following a Gaussian profile, therefore functional coupling is equal to the relative strength of excitatory connections

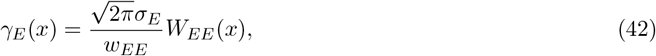

which, according to Eq. (4) is normalized such that 0 ≤ *τ*_*E*_(*x*) ≤ *τ*_*E*_(0) = 1. In Fig. 4i we show the relation between the E-E influence, obtained by inverse Fourier transform of Eq. (38), and the functional coupling, Eq. (42), for different combinations of gain values *ρ*_*E*_ and *ρ*_*I*_.

### 5.2. Network of N strongly coupled E-I pairs

In order to better understand the relationship between influence and functional coupling, we study a reduced model of *N* excitatory and *N* inhibitory rate units. Connections are all-to-all, and all connections of the same type (from population K to L, with K and L ∈ [*E, I*]) are identical. This models a strongly coupled subnetwork. In the context of our full model, for which connectivity is strictly dependent on space, this network would correspond to a population of nearby neurons, all densely connected with each other. In cortex, even if columnar, such highly coupled networks may not be as tightly organized in space as in the full model. Thus we here depart from this assumption and model a population of neurons whose functional coupling depends directly on connectivity, and not anymore on space. We construct the connectivity of this model by

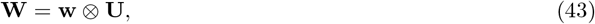

defining

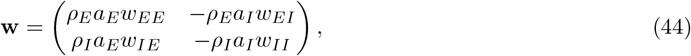

where *ρ*_*E*_ and *ρ*_*I*_ are the excitatory and inhibitory gains, respectively, the parameters 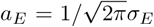 and 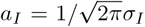 are introduced to facilitate the comparison with the full model, **U** is a *N* × *N* matrix with all elements equal to 1 and ⊗ indicates the Kronecker product. We defined the 2*N* dimensional vector **h**, the optogenetic input. Its first component is set to *h*_1_ = *δ*, and all the other i-components are set to zero, i.e. *h*_*i*_ = 0 for 1 *< i* ≤ *N*. The steady-state solution of this model

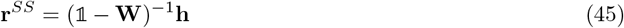

is a 2*N* -dimensional vector, where the first *N* components contain the activities of the excitatory neurons and the second *N* components the activity of the inhibitory neurons. We are interested in how stimulating neuron 1 influences the activity of the other excitatory neurons. Since the connectivity defined in Eq. (45) is uniform, all the excitatory neurons apart from the one being stimulated exhibit the same activity - which is the E-E influence. The system of Eq. (45) can be solved in closed form and, following the same notation as in Section 1.5, the influence yields

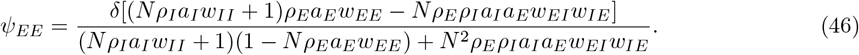

Since stability requires the denominator of this expression to be positive, this influence is suppressive if

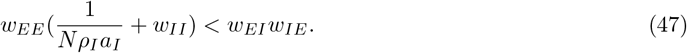

For *N* → ∞ Eq. (47) this condition does not depend anymore on the gains

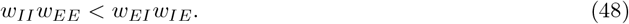

Note, however, that this depends on the normalization of the weights: if the strength of weights scale as 1*/N*, the suppression condition does not depend anymore on *N* and coincides with Eq. (41) up to a parameter *a*_*I*_. Importantly, the magnitude of influence in Eq. (46) scales as 1*/N* and thus approaches zero in the large *N* limit. Thus, the strongest influence values are expected in smaller functionally coupled subnetworks.

### 5.3. Network of 2 strongly coupled E-I pairs

Therefore, we next focus on the case *N* = 2, which consists of two pairs of E and I neurons. The first E neuron is being stimulated, while the influence is the change this stimulation induces in the second E neuron. From Eq. (46), it immediately follows that the influence for the case *N* = 2 is

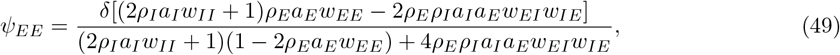

and for the same reason as above, this influence is suppressive if

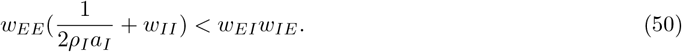

Thus, whether influence is suppressive or facilitatory in this case of two strongly coupled E-I pairs only depends on *σ*_*I*_, the gain of the inhibitory cells, but not on *σ*_*E*_, the gain of the excitatory cells. The influence as a function of these two gains is depicted in Fig. 4k, right.

### 5.4. Network of 2 E-I pairs with varying coupling strength

Finally, we study how influence between two E-I pairs depends on the coupling strength between them. We vary the coupling strength between the two pairs and compute influence as a function of the gains *σ*_*I*_ and *σ*_*E*_, to produce the plots in Fig. 4k, and l. Unlike in the full model, coupling strength is not a simple function of distance anymore, but we still assume that the cross-connections between the two pairs are symmetric, and that functional coupling is proportional to the strength of the excitatory connections between the two pairs. We further assume that inhibitory connections decrease along with the excitatory connections, when decreasing the coupling between the two pairs, and that their ratio is the same as in the full model. Concretely, if the excitatory connections between the pairs is reduced by a factor *γ*_*E*_, which takes values from 0, no-coupling, to 1, strong coupling, then the inhibitory cross-pair connections are reduced by the factor

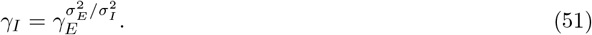

Note that, as for *γ*_*E*_ it holds that 1 ≥ *γ*_*I*_ ≥ 0, and since *σ*_*E*_ *< σ*_*I*_ the decrease of inhibitory connections is weaker than that of excitatory connections, i.e. *ρ*_*I*_ *ρ*_*E*_. This implies that there is a regime in which direct E connections approach zero while E-I-E connections are still present and provide the primary source of E-E influence. Fig. 4k left shows the dependency of influence on the gains *σ*_*I*_ and *σ*_*E*_ in this regime of small functional coupling.

Concretely, we consider a neuron model consisting of two pairs of excitatory and inhibitory neurons, whose connections are modulated by 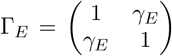and 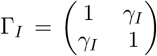, where dia gonal terms indicate the prefactors for within-pair connections, while off-diagonal terms the prefactors for the cross-pair connections.

The connectivity matrix is obtained by

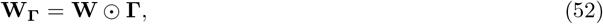

where ⊙ denotes the Hadamard product and 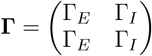. The resulting connectivity matrix is

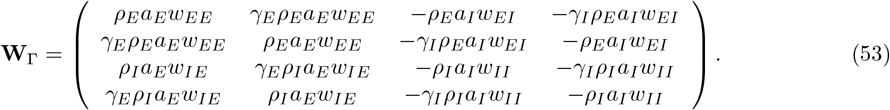

Solving Eq.(45) using the connectivity Eq.(53) and **h** as defined in the previous section we obtain the steady-state solution

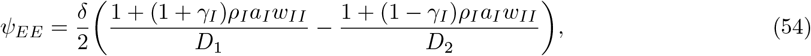

where

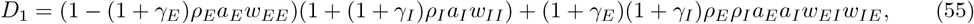

and

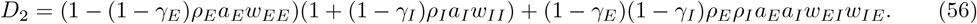

In the main text (Fig. 4k), we show Eq. (54) for *γ*_*E*_ = 0.012, *γ*_*I*_ = 0.14 (left) and for *γ*_*E*_ = *γ*_*I*_ = 1 (right) as a function of the gains *γ*_*I*_ and *γ*_*E*_. In Fig. 4l, we show Eq. (54) for fixed values of *ρ*_*I*_ and *ρ*_*E*_ as a function of the functional coupling parameter, which is equal to *τ*_*E*_. For weak coupling, the influence remains suppressive, and shows little dependence on the gains, whereas for large coupling, the influence switches from positive to negative coupling if the inhibitory gain alone, or if both gains increase. This shows that key aspects of the switching behavior observed in the experiments (Fig. 3c) can be explained by this reduced two-pair model. This enables the interpretation that switching in strongly functionally coupled subnetworks (showing high noise-correlations) happens because of strong excitatory and inhibitory reciprocal connections, whereas the absence of such switching behavior in weakly coupled (weakly noise-correlated) networks suggests weak direct excitatory connections and a dominance of disynaptic inhibition.

## 6. The I-E influence

In the experiments (Fig. 5), optogenetic stimulation is delivered to a single inhibitory neuron. As in the E-E case described above, we model this stimulation as an external input proportional to a Dirac delta centered at *x*_0_, but now targeting the inhibitory population. The Fourier transform of this input is

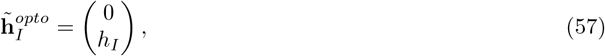

where *h*_*I*_ *>* 0 is the amplitude of the Dirac delta, and the excitatory component is set to zero, as now only the inhibitory neuron is stimulated. Given this input, we define

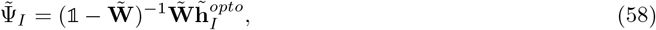

which represents the recurrent component of the steady-state solution (analogous to Eq. (12) in the E-E case).

We define the I-E influence in the Fourier space as the first component of 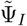, which is

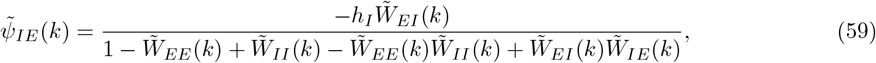

and the I-E Influence in real space is obtained by computing

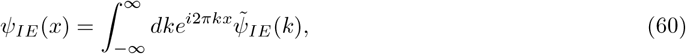

the Inverse Fourier transform of expression Eq. (59).

## 6.1. Net I-E influence

As before, we compute the net I-E influence by

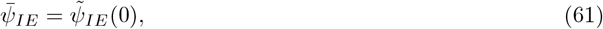

the I-E influence Eq. (59) evaluated at *k* = 0. From the normalization condition in Eq. (3), it follows that

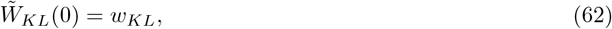

∀*K, L* ∈ [*E, I*] and thus Eq. (61) yields

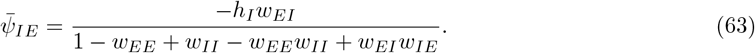

Because of the stability requirement of the network, the denominator of Eq. (63) is strictly positive. Therefore, the sign of the net I–E influence is determined solely by the numerator, which is always suppressive since *w*_*EI*_ and *h*_*I*_ are both positive by construction.

### 6.2. I-E influence depending on gain

Finally, to study how I-E influence depends on stimulus strength (contrast) we exam how Eq. (59) is modified if gains *τ*_*I*_ and *τ*_*E*_ are introduced, analogously to Section 4. For this case, we obtain Computing the spatial average, as in Eq. (63), the numerator is now

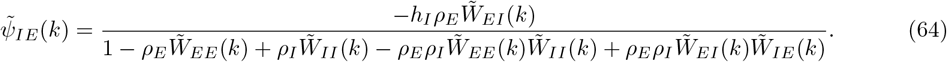

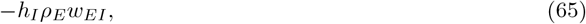

demonstrating that, in the case of I-E influence, the sign of the net influence depends only on the excitatory gain, but is unaffected by the inhibitory gain.

---

**Supplementary Figure 1.**
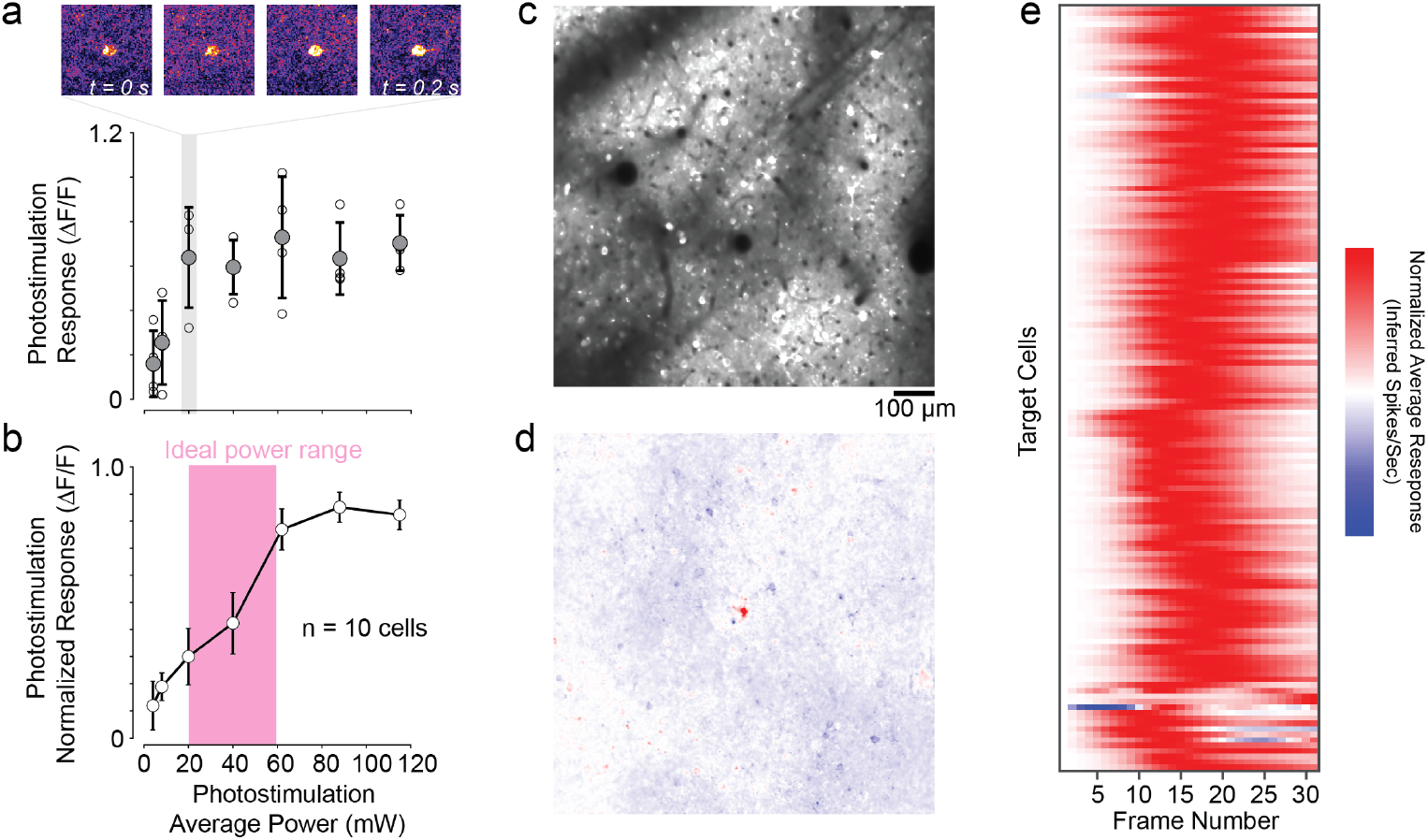
Single excitatory cell perturbations. (a) Example power response curve for a single cell. Shown are peak photostimulation responses for different average power of photostimulation laser. Shown are individual trials (open circles) and mean/SEM. (b) Example average power response curve for a single FOV. Ideal power range across experiments was 20-60 mW average power. (c) Two-photon average project of GCaMP8s single in excitatory cells. (d) Example activation of a single excitatory cell (center) during photostimulation trial. (e) Normalized average response profile across all target excitatory cells.

**Supplementary Figure 2.**
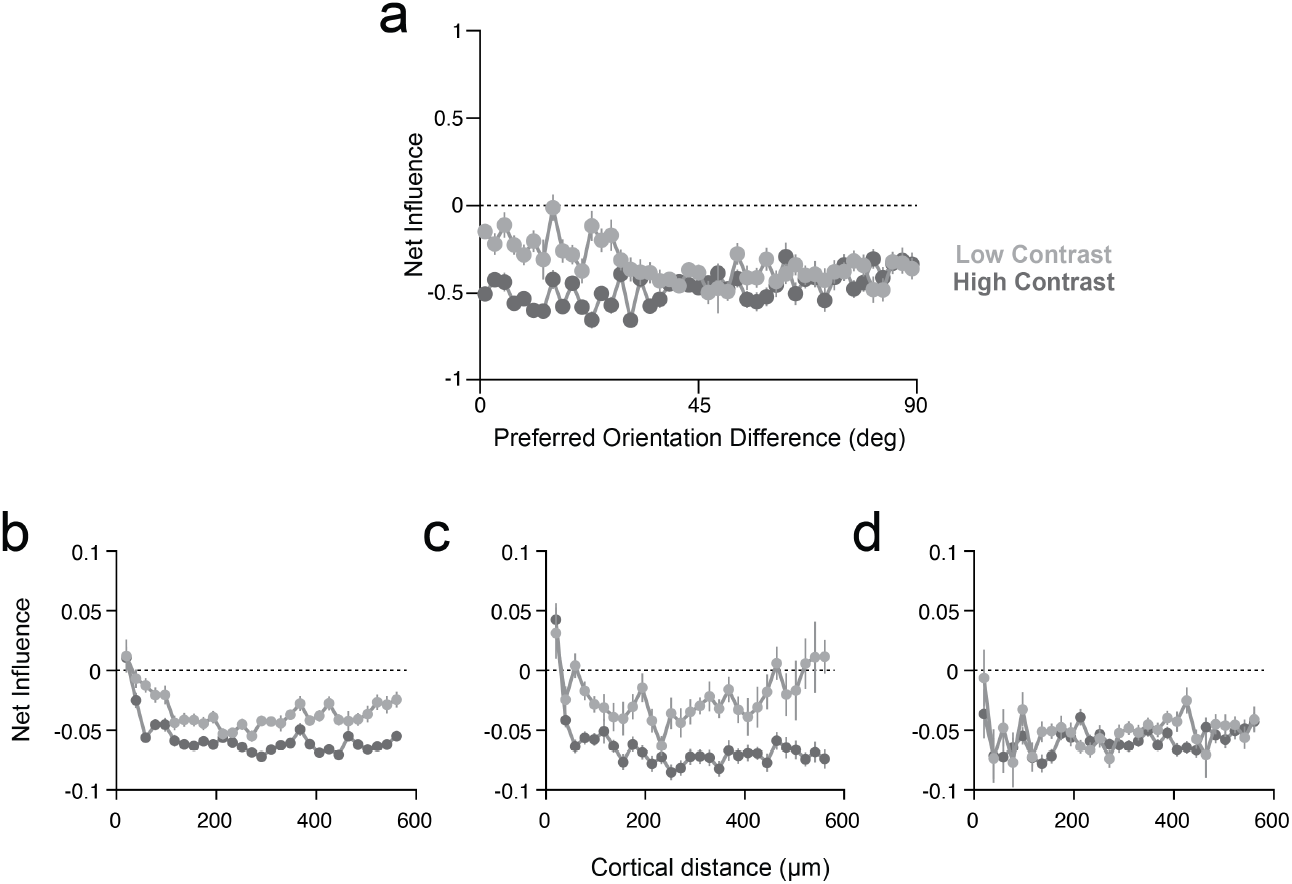
Dependence of influence on orientation similarity and contrast-dependent spatial influence profiles for nontarget cells with well-fit GLMs. (a) Relationship between difference in preferred orientation tuning and net influence for high (dark gray) and low (light gray) contrast stimuli. Influence is measured in units of change in inferred spikes per second. Distances less than 20 µm were excluded for potential photostimulation artifacts. Each data point represents mean and standard error for that distance bin. (b-d) Shown are the same plots in Figures 2c-d for nontarget cells with well-fit GLMs (average deviance <50%).

**Supplementary Figure 3.**
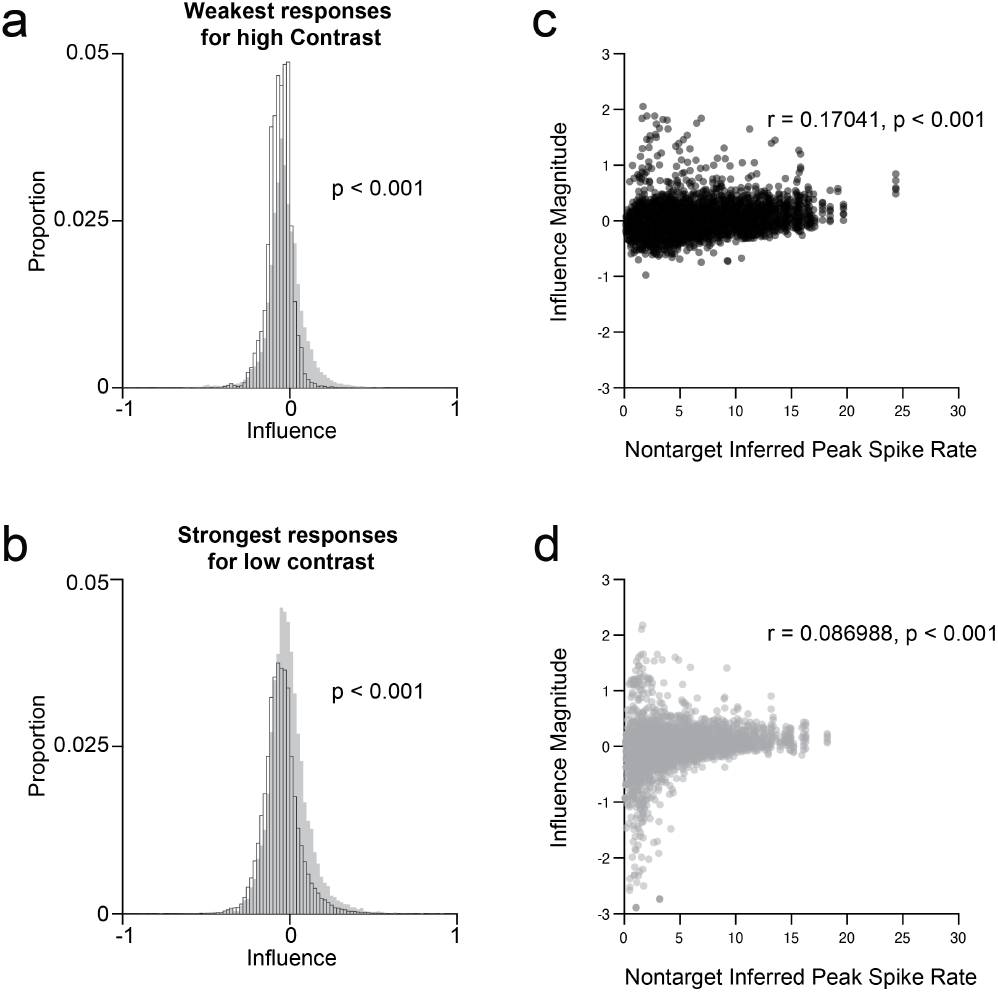
Response amplitude does not account for contrast-dependent influence shift. (a) Comparison of influences during low contrast (solid gray) with weak responding cells (bottom 25%) during high contrast (open bars). (b) Comparison of influences during high contrast (open bars) and largest responding cells (top 25%) during low contrast (solid gray). (c) Relationship between nontarget visual response amplitude during high contrast stimuli and target-nontarget influence. Each data point represents a target-nontarget pair. For each data point, the visual response amplitude is measured in the nontarget cell during sham trials. (d) Same as in **c** for low contrast stimuli.

**Supplementary Figure 4.**
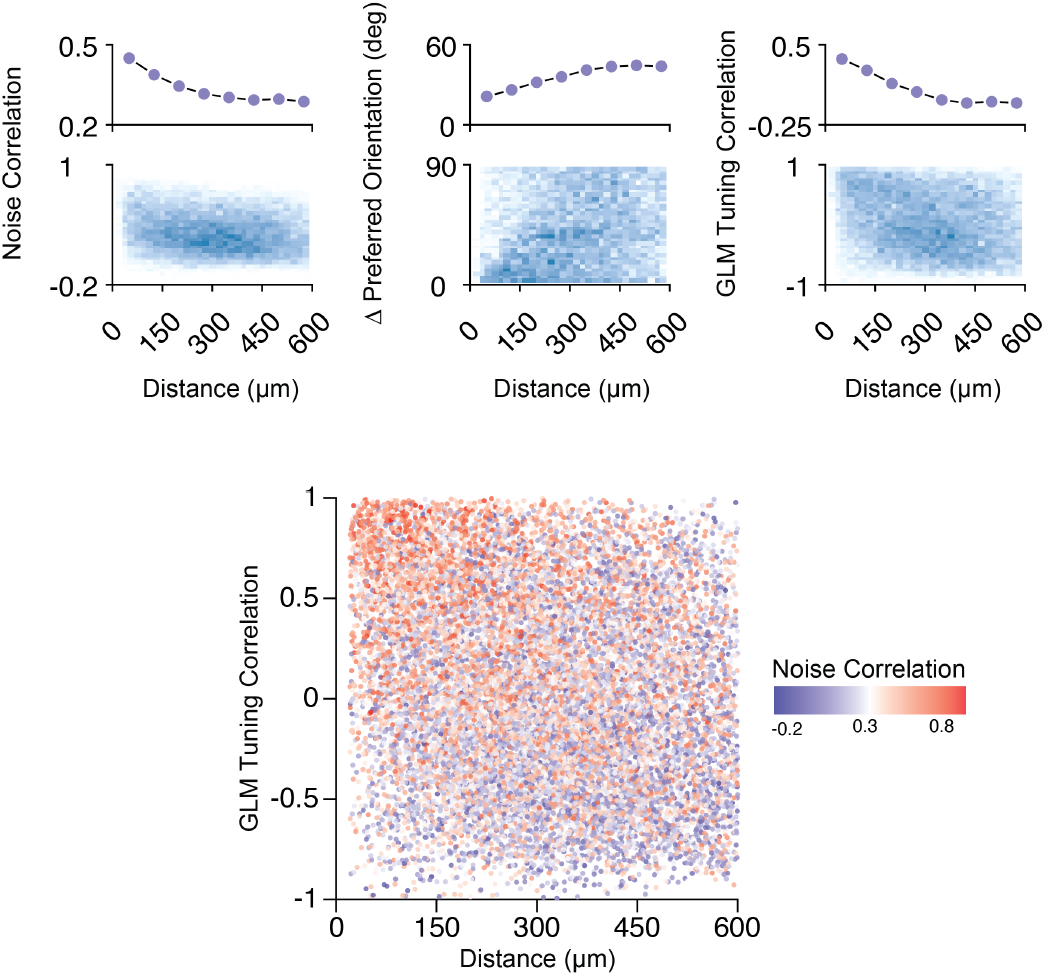
Distant-dependent functional relationships between excitatory cells in ferret V1. *Top*: noise correlation, difference in orientation preference difference, and GLM visual tuning weight correlation depend on cortical distance between excitatory cell pairs. Shown are mean values for binned distances (upper panels) and 2-D distributions (lower panels). *Bottom:* Relationship between GLM visual tuning correlation, cortical distance, and noise correlation. Each data point represents the values for a single excitatory cell pair. Nearby cells are correlated in visual tuning and have higher noise correlation. Noise correlations significantly co-vary with GLM tuning correlation (Spearman’s r = 0.25, p < 0.001, two-side test).

**Supplementary Figure 5.**
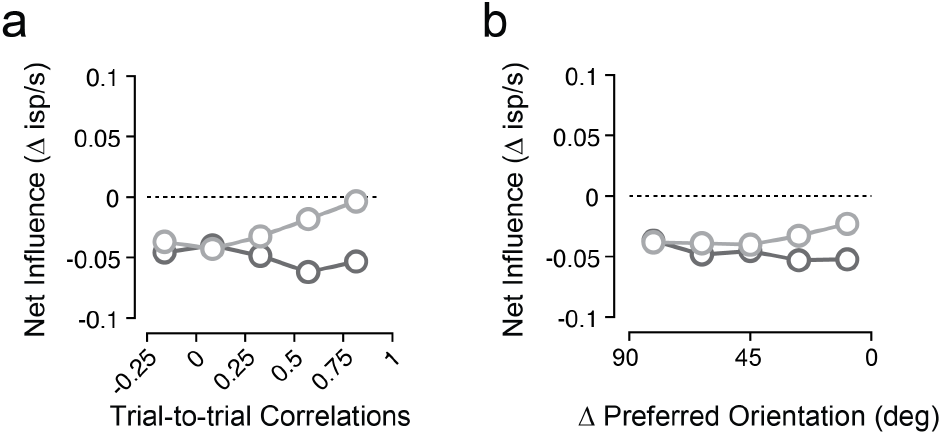
Excitatory influence dependence on trial-by-trial signal correlations and orientation preference difference. (a) Relationship between local net influence and trial-by-trial signal correlations for low (light gray) and high (dark gray) contrast. Data points are mean and standard error. Data include all perturbation trials. Spearman’s *r*_*high contrast*_ = -0.082, p < 0.001. Spearman’s *r*_*low contrast*_ = +0.15, p < 0.001. Target-nontarget pairs with high noise correlation (*r* > 0.75) exhibited more facilitatory influence at low contrast (low: mean = +0.007 ± 0.13 s.d., high: mean = -0.037 ± 0.16 s.d., p < 0.001). (b) Same as **a** for orientation preference difference. Spearman’s *r*_*high contrast*_ = -0.14, p < 0.001. Spearman’s *r*_*low contrast*_ = -0.012, p = 0.13. Target-nontarget pairs with high noise correlation (*r* > 0.75) exhibited less suppression at low contrast (low: mean = -0.037 ± 0.12 s.d., high: mean = -0.065 ± 0.12 s.d., p < 0.001).

**Supplementary Figure 6.**
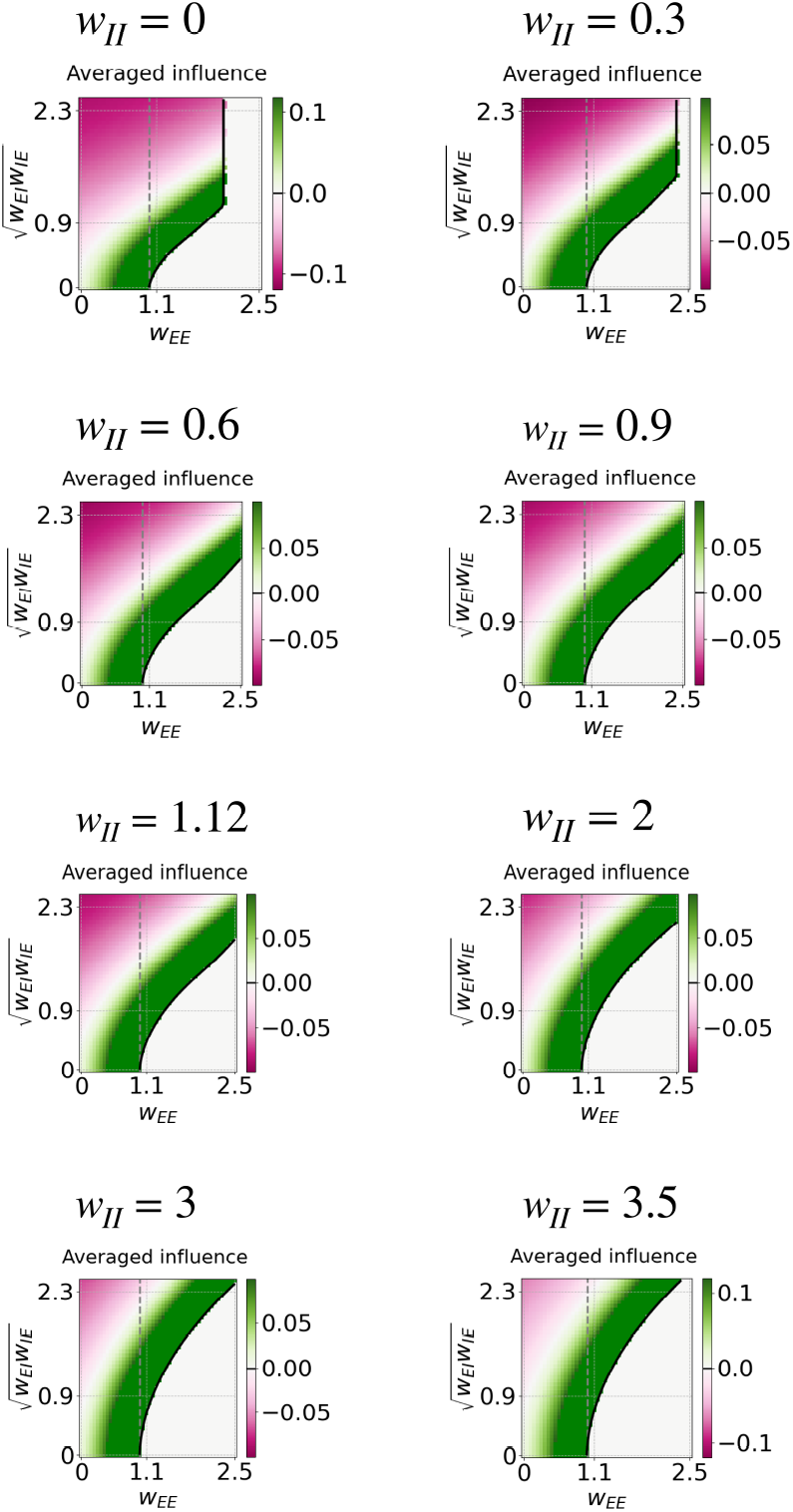
Simulated average influence with varying connection strengths between inhibitory units. Model results shown in Figure 4d reproduced for a range of *w*_*ii*_ values. Fixed parameters shown above. Overall, increased *w*_*ii*_ moves the net influence zero-crossing, but does not impact our primary finding that net suppression is observed when cross-population connection weights are large.

**Supplementary Figure 7.**
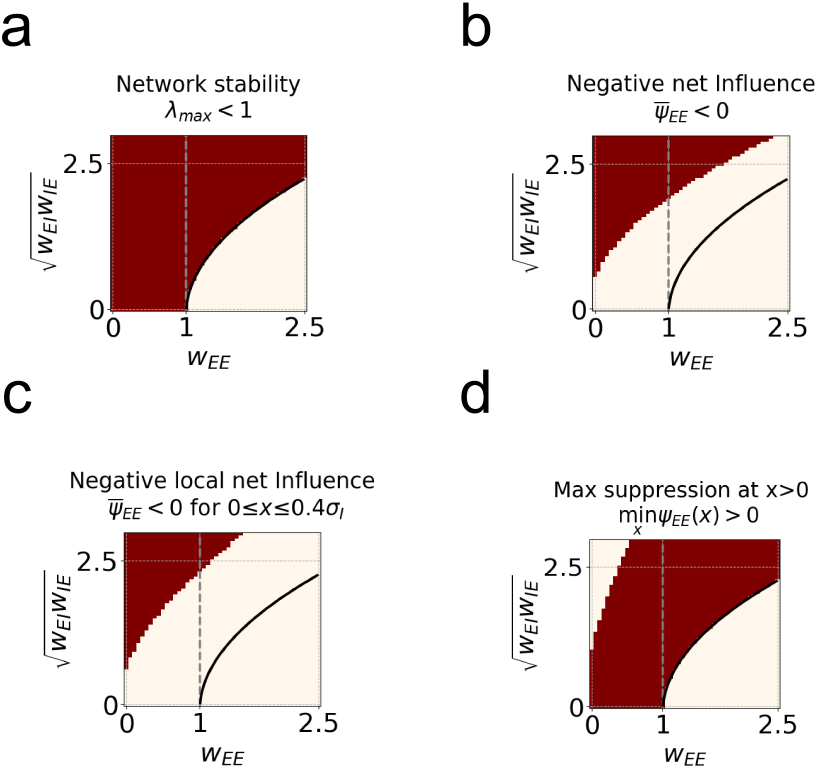
Biological plausible regime description. The area consistent with experimental observations shown in Figure 4e is defined by 4 different criteria. Here we show each criterion individually: (a) network stability, (b) averaged across space influence is suppressive, (c) locally, influence is suppressive, and (d) maximum suppression does not occur at the stimulation site.

**Supplementary Figure 8.**
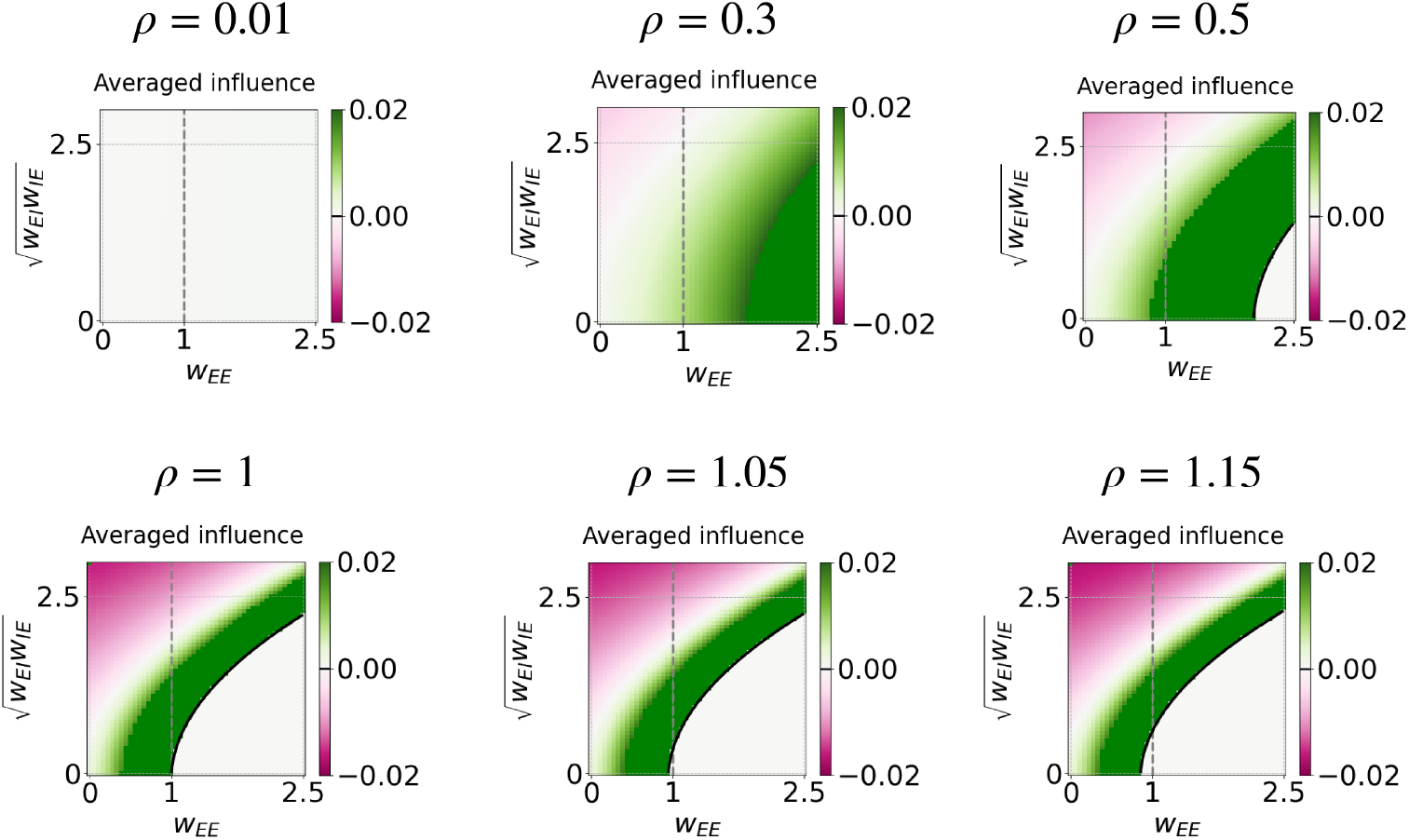
Network gain modulates spatially averaged influence across a range of connection strengths. Model results shown in Figure 4d reproduced for a range of values. For low values, the network is dominated by amplification, with a suppressive regime evolving as gain increases.

**Supplementary Figure 9.**
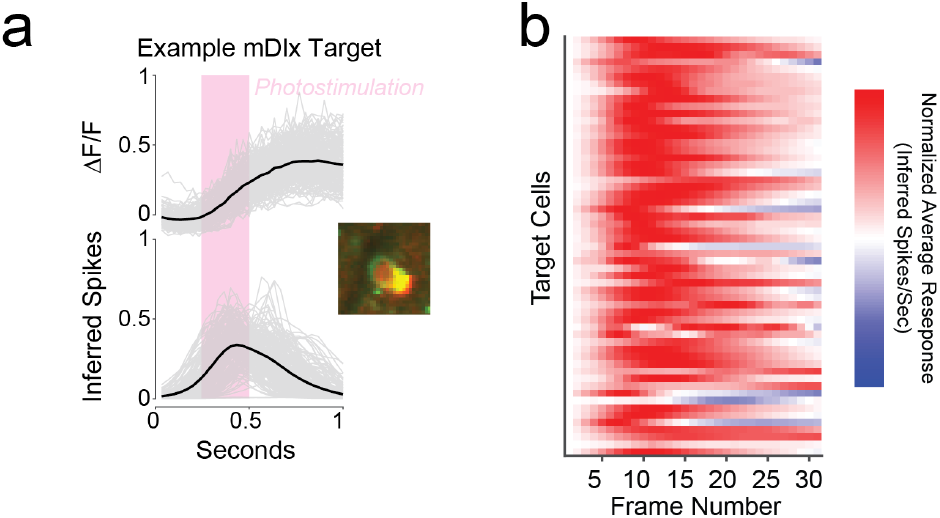
Inferring single-cell perturbation influence of targeted inhibitory interneurons onto surrounding excitatory cells. (a) Example inhibitory cell photostimulation response across trials. Data shown are individual trials (gray) and mean (black). (b) Normalized average response profile across all target inhibitory cells.

**Supplementary Figure 10.**
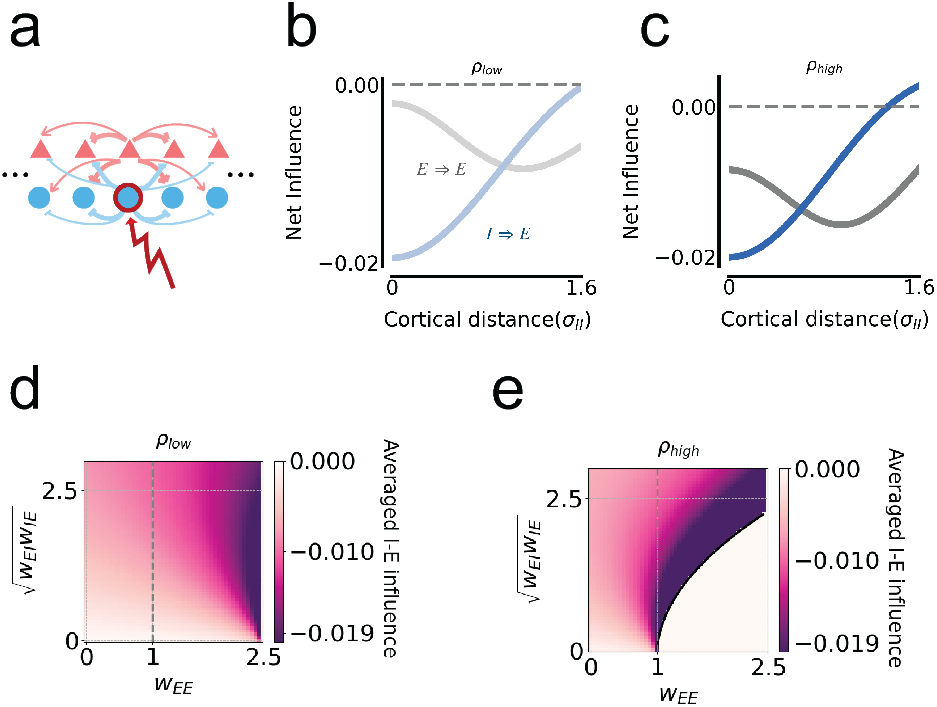
Inhibitory perturbations in Cross-Dominant model regime match experimental measurements of inhibitory-excitatory interactions. (a) Using the same model developed in Figure 4, a single inhibitory unit is perturbed. (b) Net influence for I-E (blue) and E-E (gray) interactions are shown for low gain (*ρ* = 0.4). Example uses the same model parameters as in Figure 4c. (c) Same as in b at high gain (*ρ* = 1). (d) Average I-E influence for a range of connection strengths, with the same fixed parameters as Figure 4d, at low gain. (e) Same as in d for high gain.

